# Beyond the core: endemic microbiota drive functional and microdiversity differences across salamander populations

**DOI:** 10.64898/2025.12.14.694248

**Authors:** Ostaizka Aizpurua, Elsa Brenner, Garazi Martin-Bideguren, Ion Garin-Barrio, Carlos Cabido, Antton Alberdi

## Abstract

Population-specific variation in animal microbiomes is well documented, yet the functional consequences and underlying mechanisms remain poorly understood. To address this, we conducted genome-resolved metagenomic analyses on gut and skin microbiomes from four populations of Pyrenean brook salamanders (*Calotriton asper*) inhabiting two distinct environments (Pyrenean subalpine brooks and Atlantic montane streams). From paired faecal and skin swab samples, we reconstructed 539 and 43 metagenome-assembled genomes (MAGs), respectively, and examined taxonomic composition, metabolic capacity, and microdiversity across environments. While alpha diversity remained consistent, both gut and skin microbiomes exhibited significant differences in community composition and functional potential between environments. Partitioning the gut microbiome into core, endemic, and marginal fractions revealed a dominant core community—shared across environments and accounting for over 85% of reads—that did not drive functional divergence. Instead, functional differences were primarily shaped by low-abundance, population-specific endemic bacteria. Atlantic salamanders hosted endemic taxa with significantly greater metabolic potential and higher strain-level microdiversity than those at the Pyrenees. These patterns were not explained by dietary differences and may reflect environmental influences such as temperature and nutrient availability. Our findings highlight the relevance of rare, endemic bacteria in driving microbiome function and underscore the power of genome-resolved metagenomics to uncover functional and evolutionary dynamics in wild host–microbe systems.

## Introduction

Population-specific microbiome variation is a well-documented pattern observed across a wide range of animal hosts [1], including humans [2], other vertebrates [3], and invertebrates [4]. The causes and consequences of this variation have been explored from multiple perspectives, such as industrialisation in humans [5], ecotypic differentiation in killer whales [6], and habitat degradation in howler monkeys [7]. In some cases, microbiome variation is limited to taxonomic turnover, where one bacterial species replaces another with similar functional properties, maintaining ecological stability within the host microbiome [8, 9]. Such compositional shifts often have minimal biological impact, as overall microbiome function remains unchanged [10]. In contrast, other microbiome alterations lead to significant functional differences [11], reshaping microbial dynamics and potentially influencing host physiology and health [12]. Despite the recognition of these patterns, the mechanisms driving functional divergence in microbiomes remain largely unexplored. While amphibian research has extensively documented spatial and temporal variation in skin and gut microbiome composition [13–15], the functional consequences of these taxonomic shifts remain comparatively understudied. Moreover, only a handful of studies have used metagenomic approaches to uncover the functional capacities of amphibian microbiomes [16–18].

A useful approach to understand the mechanisms driving geographical microbiota variation is to decompose its structure based on the prevalence of its members [19]. Numerous studies have attempted such classifications, often leading to varying definitions of the “core microbiome” depending on the context, study system, and research objectives [20, 21]. Within a spatial eco-evolutionary framework—where microbiomes are examined in relation to animal hosts across geographic ranges—we propose a classification into three distinct fractions: core, endemic, and marginal microbiota. Core microbiota consists of bacterial taxa consistently associated with the host across all environments and geographic regions. This group may include bacteria vertically transmitted from parents to offspring [22], as well as ubiquitous bacteria repeatedly acquired from the environment through common host behaviours, such as feeding or roosting [23]. In contrast, endemic microbiota consists of bacteria that associate with the host only in specific environments or geographic regions. Their presence may be influenced by the loss of vertical transmission, ecological factors such as environmental and dietary shifts that promote establishment of these endemic bacteria [24], or host genetic variation that exert different constraints [25]. Although not widespread across the entire host species, endemic bacteria are prevalent within their respective geographic areas, suggesting ecological processes that sustain their persistence. Endemic bacteria may introduce unique functional traits relevant to local host adaptation, such as specialised metabolic capabilities [26]. Lastly, marginal microbiota include bacteria found in only a few individuals with no consistent patterns. Such microbes may be transient, introduced through diet and environmental interactions, or more persistent but specific to individual hosts rather than population-wide trends.

To test whether microorganisms belonging to different fractions harbour different properties, we investigated the gut and skin microbiomes of Pyrenean brook salamanders (*Calotriton asper*) across two distinct environments—Pyrenean subalpine brooks and Atlantic montane streams—spanning four populations (Figure 1a-b). Using genome-resolved metagenomics, we generated the first comprehensive catalogue of bacterial genomes associated with this species and characterised the functional attributes of its microbiome. We then analysed spatial variation in functional microbiome features across environments and streams. By partitioning the microbiome into core, endemic, and marginal fractions (Figure 1c), we identified the sources of functional differences, and we employed strain-level analyses to reveal microdiversity differences across environments.

**Figure 1.**
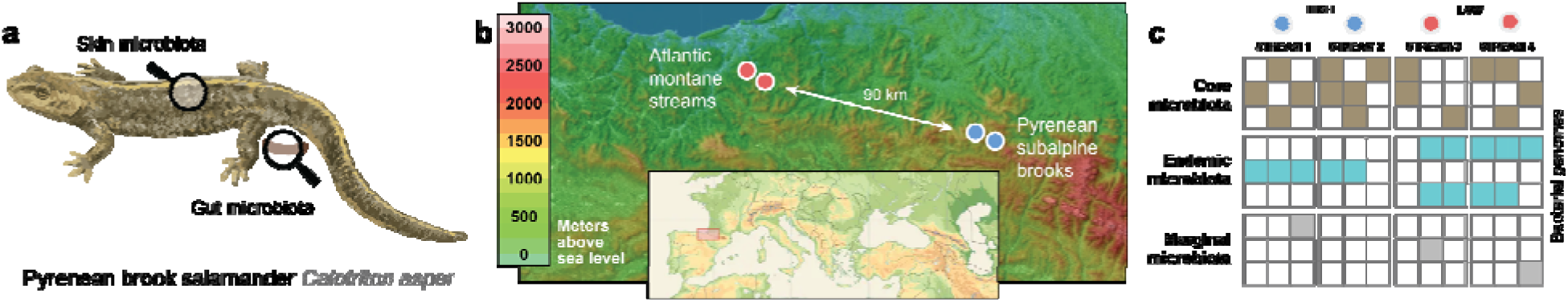
Overview of study design. **a)** Representation of the two types of samples collected from Pyrenean brook salamanders (*Calotriton asper*). **b)** Geographic location of the four streams on the western edge of the Pyrenees mountain range, coloured by environmental classification at Atlantic montane streams (red) and Pyrenean subalpine brooks (blue). **c)** Diagram of the theoretical partitioning of the gut microbiota into three fractions: core, endemic and marginal.

## Materials and methods

This study was conducted within the framework of the Earth Hologenome Initiative (EHI), thus adhering to its standardised methodological procedures for sample collection, preservation, laboratory processing and bioinformatics [27].

### Study species

The Pyrenean brook salamander (*Calotriton asper*) is an endemic species to the Pyrenees, the 800 km-long mountain range that separates the Iberian Peninsula from the rest of continental Europe (Figure 1a). Along their distribution range, populations display distinct genetic diversity with a positive correlation with altitude [28]. Surrounded by peaks exceeding 3,400 m above sea level (ASL), these salamanders predominantly inhabit high-elevation (up to 2,500 m ASL) streams and lakes with sparse vegetation, where they feed primarily on aquatic invertebrates.

While the main population within the Pyrenean mountain range is abundant and well-connected, smaller, peripheral low-elevation populations exist in the surrounding mountains. One such population resides close to the Atlantic coast in the Basque Country, approximately 90 km from the main population. These isolated peripheral populations inhabit markedly different environments compared to the Pyrenean populations, living in spring-fed streams surrounded by dense Atlantic forests. Given these environmental contrasts, it is likely that the two populations have developed different adaptations to thrive under their respective environmental conditions.

### Field sampling

Field sampling was conducted in four streams in August 2023 in the Northern Iberian Peninsula. The two Pyrenean subalpine brooks, Erlan and Harpea, are situated in different watersheds on the eastern edge of the Pyrenees mountain range at elevations exceeding 1,000 meters ASL, separated by an approximate linear distance of 3 km. In contrast, the two Atlantic montane streams, Goizueta and Leitzaran, are located approximately 90 km west of the Pyrenees’ western edge, at elevations less than 500 meters ASL, separated by an approximate linear distance of 4 km (Figure 1b).

31 adult male newts (Pyrenean subalpine streams = 15 and Atlantic montane streams = 16) were captured with hand-nets, weighed, measured, and kept in individual containers containing stream water until defecation or for up to two hours. Animal handling was conducted using nitrile gloves, replaced between each animal, ensuring no direct contact between the newt and human skin. Two types of samples were collected from each captured individual: skin swabs and faecal samples. Skin samples were obtained following the recommended amphibian swabbing protocols described in Hyatt et al. [29] and Boyle et al. [30] modified for microbial analysis by first rinsing the newt with sterile water to reduce debris and wash away environmental microbes, and then gently swabbing the dorsal and ventral skin for 30 seconds each, using single-use sterile FLOQSwabs (Copan Diagnostics, Italy) moistened with sterile water. Faecal samples were collected from the water immediately after defecation. Both faecal samples and skin swabs were stored in standardised EHI collection tubes, containing 1 mL DNA Shield (Zymo, cat. Number R1100-250). Samples were frozen within the day and kept at −20°C until DNA extraction.

### Laboratory sample processing

Samples were mechanically lysed using Lysing Matrix E 96-tube Rack (1,2 mL, MP Biomedicals™, cat. Number 116984010) on a TissueLyser II (Qiagen) for 2 times 6 minutes at 30 Hz, inverted between TissueLyser runs. In the case of skin samples, swabs were kept inside the tube to maximise cell and DNA elution into the buffer. Subsequently, samples were centrifuged and DNA extraction was conducted from 200 μl of the supernatant using the standardised EHI DNA extraction procedure DREX [31]. Samples were randomised to minimise batch effects, and DNA extraction blanks were included at the beginning of the extraction. These blanks consisted of the same preservation buffer that was used for sample storage and were included throughout all laboratory steps.

DNA sequencing libraries were generated using the Blunt-End Single Tube (BEST) protocol with BEDC3 adaptors [32]. Libraries were purified by adding 1.67 times the library volume of Solid Phase Reversible Immobilisation (SPRI) beads. The mixture was incubated for 5 min at room temperature (ca. 21°C), followed by a double wash with 80% ethanol on a 96S Super Magnet (Alpaca, SKU: A001322). The purified DNA was eluted in Elution Buffer Tween (EBT; Buffer EB, Qiagen, cat. Number 19086, and TWEEN® 20, Sigma-Aldrich, cat. Number P9416-50ML) solution by incubating at 37°C for 10 mins. Library preparation performance was assessed by means of a qPCR assay conducted on a Mx3005 qPCR System (Agilent, USA) using 1:20 diluted libraries. qPCR mixes were composed of 2.5 µL 10x PCR Gold Taq buffer, 2.5 µL MgCl_2_ (25 mM), 0.2 µL dNTP mix (10 mM each), 1 µL (10 µM) of IS7 and IS8 primers, 0.5 µL AmpliTaq GOLD DNA polymerase, 1 µL SYBR green dye, 2 µL diluted sample library and 14.3 µL sterile dH_2_O for a total reaction volume of 25 µL. Amplification was carried out following the program: 1) 1 cycle of 95°C for 12 mins, 2) 40 cycles of a) 95°C for 20 sec, b) 60°C for 30 sec, and c) 72°C for 40 sec, 3) dissociation curve. Amplification curves from qPCR were used to decide the optimal number of cycles for each library to reach the PCR amplification plateau, yet without overamplifying the libraries to avoid excessive clonality.

Subsequently, sequencing libraries were PCR-amplified using unique dual-index Illumina primers per sample. PCR reaction was 5 µL 10x PCR Gold Taq buffer, 5 µL MgCl_2_ (25 mM), 0.4 µL dNTP mix (10 mM each), 1 µL (10 µM) for each P7 and P5 primers, 1 µL AmpliTaq GOLD DNA polymerase, 26.6 µL sterile dH_2_O, and 10 µL sample library for a total reaction volume of 50 µL. The PCR reaction was 1) 1 cycle of 95°C for 12 mins, 2) 7-20 cycles of a) 95°C for 20 sec, b) 60°C for 30 sec, and c) 72°C for 40 sec, 3) 1 cycle of 72°C for 5 min, and 4) hold at 4°C. Each sample was amplified for a different number of cycles (7-20), as determined by the previous qPCR assay.

The indexed libraries were purified with SPRI beads using 1:2 bead-to-library ratio. The mixture was incubated for 5 min at room temperature (ca. 21°C), followed by a double wash with 80% ethanol on a 96S Super Magnet (Alpaca, SKU: A001322). The purified DNA was eluted in EBT solution by incubating at 37°C for 10 min. Purified indexed libraries were pooled together and sequenced on an Illumina NovaSeqX platform, aiming for 5 GB (approximately 17 million reads) of data per sample.

### Bioinformatic data processing

Raw sequencing data were processed using the standard EHI bioinformatic pipeline [33]. Briefly, reads were quality trimmed using fastp [34] before being mapped using Bowtie2 [35] to the *Calotriton arnoldi* reference genome (GCA_963921515.1), which was the closest reference genome to *Calotriton asper* available at the time of the analysis. Host reads were subsequently filtered from BAM files using samtools [36]. Non-host reads were then assembled both individually and coassembled per stream using MEGAHIT [37]. Single-coverage or multi-coverage binning was then performed using CONCOCT [38], Maxbin2 [39] and MetaBAT2 [40], before being refined using MetaWRAP’s bin_refinement module [41, 42]. MAGs were dereplicated at 98% ANI using dRep [43] with MASH [44] and FastANI [45], and non-host reads were mapped to the dereplicated MAG catalogue using Bowtie2, with coverage statistics calculated using CoverM [46].

MAGs were quality-checked with CheckM2 [47], and annotated taxonomically using GTDBtk [48] with the GTDB r214 [49]. The phylogenetic tree of MAGs was generated through pruning the reference GTDB genomes used for phylogenetic placement. DRAM [50] was used to annotate MAG functions using multiple reference databases [51–58]. Subsequently, gene annotations were distilled into Genome-Inferred Functional Traits (GIFTs) using distillR [59], producing biologically meaningful annotations that highlight each bacterial genome’s potential to degrade or synthesise compounds relevant to host metabolism. DistillR utilises a curated database of over 300 metabolic pathways, employing KEGG and Enzyme Commission (EC) identifiers to calculate standardised GIFT values (Table S1). These values range from 0 to 1, where 0 signifies the absence of all genes associated with a particular metabolic pathway, and 1 indicates the presence of all necessary genes. For example, if a pathway step requires two specific identifiers, it is deemed complete when both are present, half-complete when only one is present, and empty if neither is present. In the case of single-locus traits, the GIFT becomes binary. The domains considered encompass functions relevant for both gut (e.g., polysaccharide degradation, vitamin biosynthesis) and skin (e.g., toxin biosynthesis) microbiomes.

The fraction of bacterial and archaeal DNA in our samples was estimated using SingleM’s prokaryotic_fraction function [60, 61]. SingleM estimates the expected genomic data from prokaryotic origin in a sample by measuring the coverage of prokaryotic single-copy marker genes, and inferring genome sizes based on the closest relatives present in public databases. The Domain-Adjusted Mapping Rate (DAMR) was calculated as the rate of read mapping to the MAG catalogue divided by the fraction of the community predicted to be bacterial or archaeal.

Dietary analysis was conducted using BarcodeMapper [62], which quantifies plant, fungal, and animal DNA from shotgun sequencing datasets by mapping reads to taxonomically annotated COI and ITS marker gene sequences. Pairwise microdiversity dissimilarity metrics were computed following the population-ani method implemented in lorikeet [63].

Genomic differences between animals from Atlantic montane streams and Pyrenean subalpine brooks were analysed using kmer-based genome skimming, due to the impossibility of generating sufficient data for variant calling based on the large size of *Calotriton* genomes.

Using a custom-made pipeline (https://github.com/alberdilab/genome_skimming), we extracted the sequencing reads mapping to *Calotriton arnoldi* and analysed them using skmer [64], which enables computing distances between sequence read pools based on their combined kmer profiles. After filtering out samples with fewer than 5 million reads mapped to the reference genome to reduce noise, we tested for significant differences between the contrasted populations using PERMANOVA.

### MAGs diversity and statistical analysis

All statistical analyses were conducted using R software v.4.3.2 [65], and were compiled as a Rmarkdown website enabling full reproduction. We calculated the alpha diversities of microbial communities using Hill numbers [66]. To capture the effects of different diversity components (richness, neutral, phylogenetic) and diversity orders (q = 0 considers only presence/absence, while q = 1 gives weight to MAGs based on their relative abundances), we calculated species richness at q = 0, neutral diversity at q = 1, phylogenetic diversity at q = 1 using the package Hilldiv2 [67]. Alpha diversity differences were determined by a parametric t-test when the data were normally distributed and when the variances between contrasted groups were equal. When these assumptions were not held, a non-parametric Wilcoxon test was performed.

We calculated compositional dissimilarities across samples using Hill numbers by computing the Jaccard-type turnover for neutral and phylogenetic beta diversities at order q = 1 using hilldiv::hillpair. To visualise the variation in microbial composition, we performed nonmetric multidimensional scaling (NMDS) ordination plots based on the derived distance matrices. Differences in dispersion within sampling methods were assessed using the betadisper function in the vegan package [68]. To test for differences in microbial composition between samples, we conducted a PERMANOVA using the adonis2 function in vegan and pairwise comparisons using pairwiseAdonis [69]. Microbial differential abundance analysis was performed using ANCOM-BC2 [70]. Additionally, to visualise bacteria according to their functional traits, MAGs were ordinated based on their GIFTs (Genomic Information Functional Traits) through PCoA[71]. In this ordination, MAGs with similar functional traits are displayed closer to each other while MAGs with distinct profiles are placed further away. To analyse functional differences, community-weighted averages of GIFTs were calculated using distillR::to.community, which weights each MAG’s traits by its abundance in the sample. Differences between environments were calculated using the Kruskal-Wallis test, followed by the Bonferroni–Holm method for multiple comparisons to obtain adjusted p-values.

To assess the influence of bacteria with different levels of prevalence, we partitioned MAGs into three fractions. Genomes detected in all four streams were considered core microbes. Genomes detected in both streams within the same environment, with a minimum prevalence of 20%, were classified as endemic bacteria. Genomes detected at only one environment and with less than 20% prevalence were considered marginal.

## Results

We captured 31 adult male Pyrenean brook salamanders (*Calotriton asper*) from two Pyrenean subalpine brooks (n=8+7) and Atlantic montane streams (n=7+9). Individuals from Pyrenean subalpine brooks exhibited significantly greater snout–vent lengths (t-test: t = 2.09, p-value = 0.0046). However, no significant differences in weight were detected between environments (t-test: t = 0.82, p-value = 0.419). Given the current limitations to reliably characterise and validate the representativeness of other members of the microbiome (e.g., fungi, viruses), we limited our analysis to bacterial and archaeal components.

### Bacterial genome catalogues associated with *Calotriton asper*

We produced a total of 154.38 and 162.04 Gb (gigabases) of metagenomic data from faecal samples and skin swabs to characterise the gut and skin microbiomes of salamanders. From these data, we generated two reference bacterial genome catalogues (gut and skin) with significantly different features (Figure 2a). The gut genome catalogue included 539 metagenome-assembled genomes (MAGs), with a mean completeness of 85.01 ± 16.2% and a contamination level of 1.68 ± 2.25% (Figure S1). An average of 3.66 ±1.24 Gb out of the 4.98 ± 1.49 Gb sequenced per sample was mapped against the reference catalogue (Figure S2), yielding a near-complete (99.94 ± 0.24%) estimated recovery of bacterial and archaeal genomes (Figure S3). In contrast, despite a nearly identical sequencing effort, the skin genome catalogue only recovered 43 genomes. The reconstructed MAGs had a mean completeness of 77.24 ± 17.04% and a contamination level of 4.48 ± 2.42% (Figure S4). Only 0.25 ± 0.38 Gb out of the 4.47 ± 2.21 Gb of data sequenced per sample mapped against the genome catalogue (Figure S5), with an estimated microbial fraction recovery of 54.18 ± 12.63% (Figure S6).

**Figure 2.**
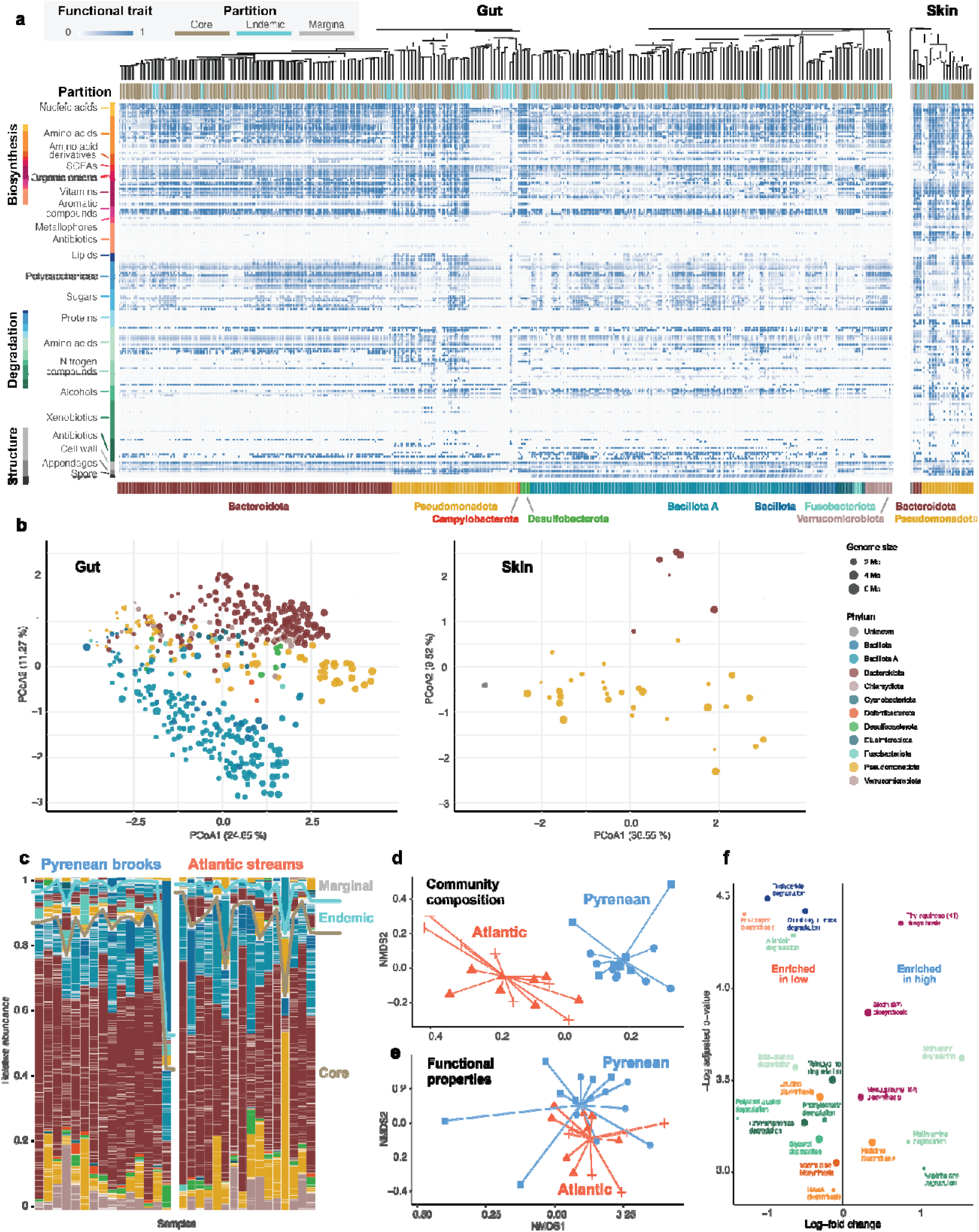
Composition and variation of the gut and skin microbiomes. **a)** Phylogeny, partition and functional characterisation of microbial genomes reconstructed from faecal (gut) and skin samples. Each tile represents a metabolic function, with lighter shades indicating lower capacity to accomplish it and darker shades indicating higher capacity. **b)** Two-dimensional PCoA ordinations of gut and skin MAGs based on their functional traits. **c)** Relative abundances of gut microbiomes in animals from Pyrenean subalpine brooks and Atlantic montane streams, sorted by MAG partitioning into core, endemic and marginal fractions. **d)** Compositional dissimilarity of the gut microbiome across animals from Pyrenean subalpine brooks (blue) and Atlantic montane streams (red). Shapes indicate different streams. **e)** Functional dissimilarity of the gut microbiome across animals from Pyrenean subalpine brooks (blue) and Atlantic montane streams (red). Shapes indicate different streams. **f)** Enrichment of gut microbiome’s metabolic functions in samples from Pyrenean subalpine brooks (left) and Atlantic montane streams (right).

The gut genome catalogue comprised 13 bacterial phyla, with Bacteroidota (56.02 ± 16.86%), Bacillota A (17.72 ± 6.39%), and Pseudomonadota (11.57 ± 13.50%) being the most abundant, collectively accounting for 84.05% of MAGs (Figure S7). Among the 539 MAGs, 91.65% lacked species-level annotation, and 24.6% lacked genus-level annotation. Species-level annotations were observed in only three phyla: Pseudomonadota (48.28% of MAGs with species-level annotation), Fusobacteriota (33.33%), and Bacillota (4.35%), while the rest of the recovered genomes were new to science. The dereplicated MAG catalogue contained 1,219,667 redundant genes, of which 942,120 (77.24%) were annotated and 557,611 (59.19%) were assigned KEGG orthologs. The functional ordination exhibited a huge phylogenetic signal: genomes belonging to the same phyla aggregated together, though with significant dispersion within the main clades (Figure 2b).

The skin genome catalogue comprised bacteria from only two phyla: Pseudomonadota—35 MAGs accounting for 84.9 ± 15.8% of the sequencing reads—and Bacteroidota—6 MAGs capturing 14.9 ± 15.8% (Figure S8). Among the 43 MAGs, 76.74% lacked species-level annotation, and 11.63% lacked genus-level annotation. The MAG catalogue contained 114,303 redundant genes, of which 88,006 (76.99%) were annotated and 52,991 (60.21%) were assigned KEGG orthologs. The functional ordination showed that skin bacteria encompassed a narrower functional landscape compared to gut bacteria, despite the large functional variability within Pseudomonadota (Figure 2b). Two genomes with very low completeness failed to get phylum-level annotation, but clustered with Pseudomonadota in the functional ordination, suggesting functional similarity despite the lack of taxonomic information.

### Microbiota differences between environments

The gut microbiota of salamanders exhibited comparable alpha diversity values across environments (LM: t-value 7.194, p-value > 0.05, Figure S9). However, beta diversity analyses revealed distinct community compositions between environments (PERMANOVA-neutral: R^2^ = 0.234, p = 0.001; Figure 2d). A total of 180 MAGs were differentially abundant between environments, yet without consistent enrichment patterns of major microbial lineages. While the average metabolic capacities of the bacterial communities were nearly identical in both environments (Wilcoxon: W = 149, p-value = 0.137), the functional properties of microbiomes differed significantly (PERMANOVA: R^2^ = 0.90, p-value = 0.022; Figure 2e). Gut microbiomes of Atlantic animals showed significantly higher biosynthetic and degradation capacities of multiple metabolites recognised as relevant for hosts, such as leucine, allantoin or phenylacetate (Figure 2f).

To determine whether diet contributed to the observed faecal microbiome differences, we examined the non-bacterial, non-host fraction of our metagenomic data, which contained remnants of invertebrates, vertebrates, plants, and fungi— all potential contributors to gut community composition via direct consumption or secondary ingestion (Figure S10). However, this snapshot did not reveal any significant variation in host diet between environments (Wilcoxon: W = 101.5, p = 0.475; Figure S11). Similarly, the genome skimming analysis also failed to reveal significant genetic differences between animals from different environments (PERMANOVA: R^2^ = 0.05, F=0.915, p-value=0.715).

The skin microbiota of salamanders from Pyrenean subalpine brooks displayed increased richness compared to their Atlantic counterparts (Wilcoxon: p = 0.007), while neutral and phylogenetic alpha diversities remained consistent across elevations (Figure S12). Beta diversity analyses revealed distinct community compositions, with stream-specific effects observed in the Atlantic montane streams (PERMANOVA-neutral: elevation, R² = 0.151, p = 0.001; stream, R² = 0.111, p = 0.018). Accordingly, 10 MAGs were differentially abundant between Pyrenean subalpine brooks and Atlantic montane streams. Functional trait analysis revealed significant differences between the microbiota of salamanders from both environments. The overall metabolic capacity was higher in Atlantic montane streams (0.385 ± 0.0503) compared to animals from Pyrenean subalpine brooks (0.326 ± 0.0525) (Wilcoxon: W = 183, p < 0.001), with multiple metabolic functions showing environment-specific patterns (Figure S13).

### Contribution of microbiota fractions to functional differences

To explore the sources of functional capacity differences, we partitioned the microbial communities into core, endemic, and marginal fractions, according to the prevalence of each genome across streams and elevations. For our analyses, the “core” microbiota referred to MAGs present in samples from all four streams. This core encompassed 389 MAGs (72.2%) (Figure 3a). The endemic fraction—MAGs present at only one environment— contained 99 genomes (18.3%). Finally, the marginal fraction contained bacteria sparsely represented in only a few individuals; it consisted of 51 MAGs (9.5%). In terms of relative abundances, the averaged aggregated representation of core bacteria across all animals was 87%, followed by 8.8% of endemic and 4.1% of marginal bacteria (Figure 3b). The average genome size (Wilcoxon: W = 21740, p.adj < 0.001; Figure 3c) and metabolic capacity (Wilcoxon: W = 21482, p.adj < 0.001; Figure 3d) of core bacteria were significantly higher than those belonging to the endemic and marginal fractions. Functional capacity differences between core and endemic bacteria were mainly driven by greater genomic potential for organic anion and vitamin biosynthesis, as well as degradation of polysaccharides and amino acids. In addition, the functional attributes of Pyrenean and Atlantic endemic bacteria also differed significantly, with bacteria endemic to Atlantic montane streams displaying higher overall metabolic capacities (Wilcoxon: W = 552, p < 0.002), with increased capacities for a number of functions, including amino acid biosynthesis, sugar degradation and antibiotic degradation.

**Figure 3.**
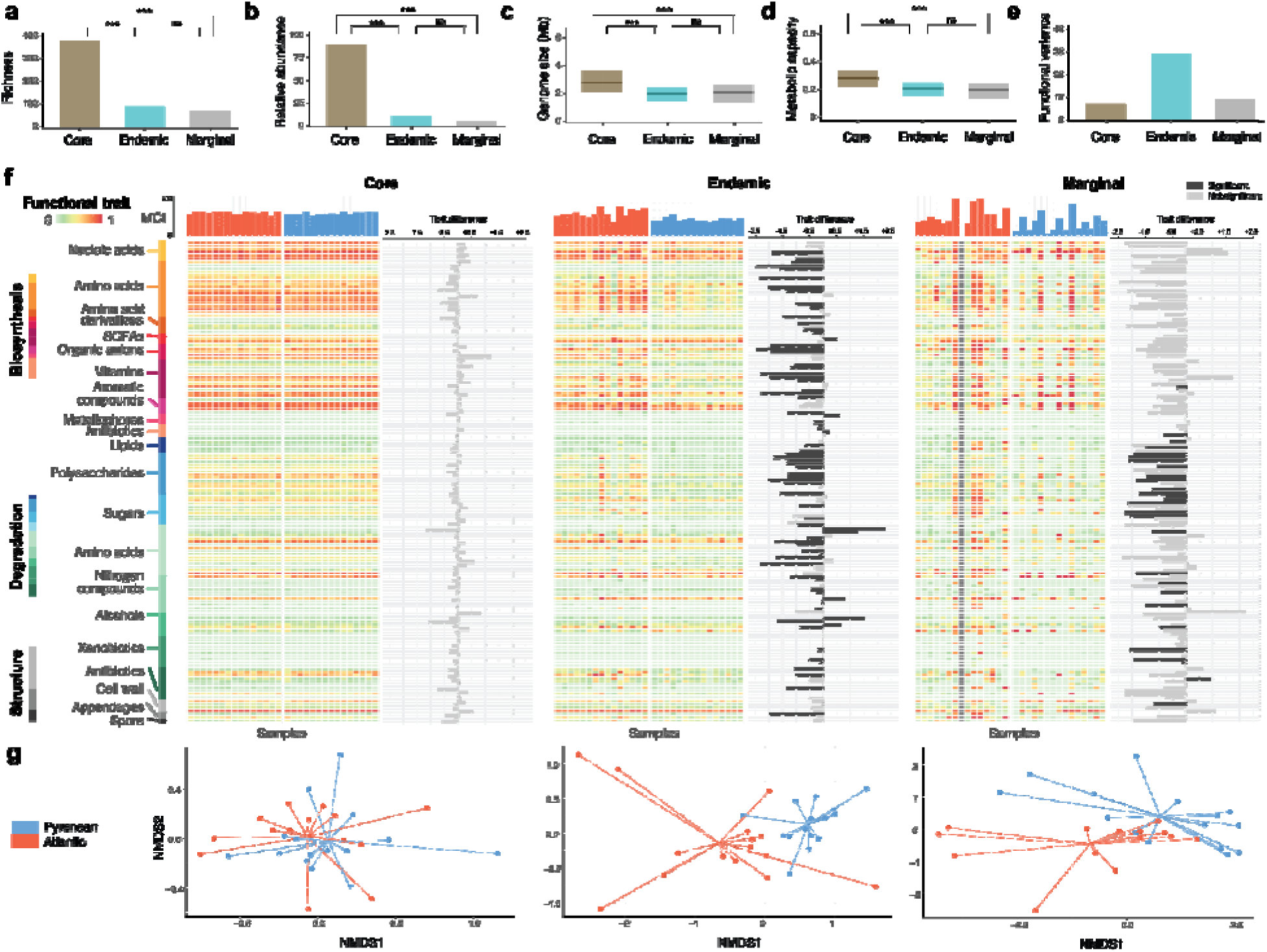
Microbiome partitioning into core, endemic and marginal fractions. **a)** Number of genomes assigned to the different fractions. **b)** Average cumulative relative abundance of the genomes. **c)** Average size of the genomes. **d)** Average metabolic capacity of the genomes. **e)** Percentage of functional variation explained by the genomes. **f)** CommunitylJweighted averages of genome-inferred metabolic traits, partitioned by core, endemic and marginal fractions.

These differences between core and endemic bacteria, along with variations in endemisms, influenced how bacteria contribute to the functional differences between communities in animals from different environments (Figure 3f). Core and marginal bacteria both exhibited differences between environments (p = 0.003 and p = 0.014, respectively), though the variance explained was modest (11.9% and 11.1%). No GIFTs differed significantly between environments in the core microbiota, whereas 34 GIFTs did so in the marginal fraction (Figure 3e). The endemic fraction demonstrated the largest effect size, with an R² of 0.29 and 89 GIFTs with differing values between Pyrenean subalpine brooks and Atlantic montane streams, resulting in a highly significant difference between environments (p < 0.001) (Figure 3g).

In the case of the skin, the core microbiome encompassed 25 MAGs (58%), while the endemic fraction contained 7 (16%) and the marginal fraction 11 (25.6%). Considered together, the core bacteria represented 90% of aggregate relative abundance, followed by 13.2% of endemic and 8.7% of marginal bacteria. The low number of genomes per category precluded us from conducting more detailed analyses.

Heatmaps display mean trait values, with darker shades of red indicating greater capacity to carry out the specific function; bar plots above each heatmap show the overall metabolic capacity index (MCI) per sample (the mean across all traits) in Atlantic montane streams (red) and Pyrenean subalpine brooks (blue). Horizontal bar plots indicate the difference between mean values in each environment, with dark colours indicating statistically significant differences between environments. **g)** NMDS ordinations of the samples ordinated according to the community-weighed functional properties of core, endemic and marginal microbiota fractions.

### Microdiversity differences between environments

To assess strain-level variability among gut bacteria, we compared reconstructed genomes across environments and microbiome fractions. Although both environments harboured the same core bacterial species, strains isolated within the same environment exhibited higher average nucleotide identity (ANI) to one another than to strains from the other environment, highlighting microdiversity differences between environments (Figure 4a). Moreover, strains from Atlantic montane streams showed significantly greater microdiversity than their Pyrenean counterparts (Wilcoxon: W = 8720, p-value < 0.001; Figure 4b). A similar pattern was observed among endemic taxa: microdiversity values of Atlantic endemics exceeded those of Pyrenean endemics (W = 70662, p-value < 0.001; Figure 4c). Together, these results indicate that—despite comparable overall gut-microbiome diversity across environments—within-species microdiversity was greater at Atlantic montane streams.

**Figure 4.**
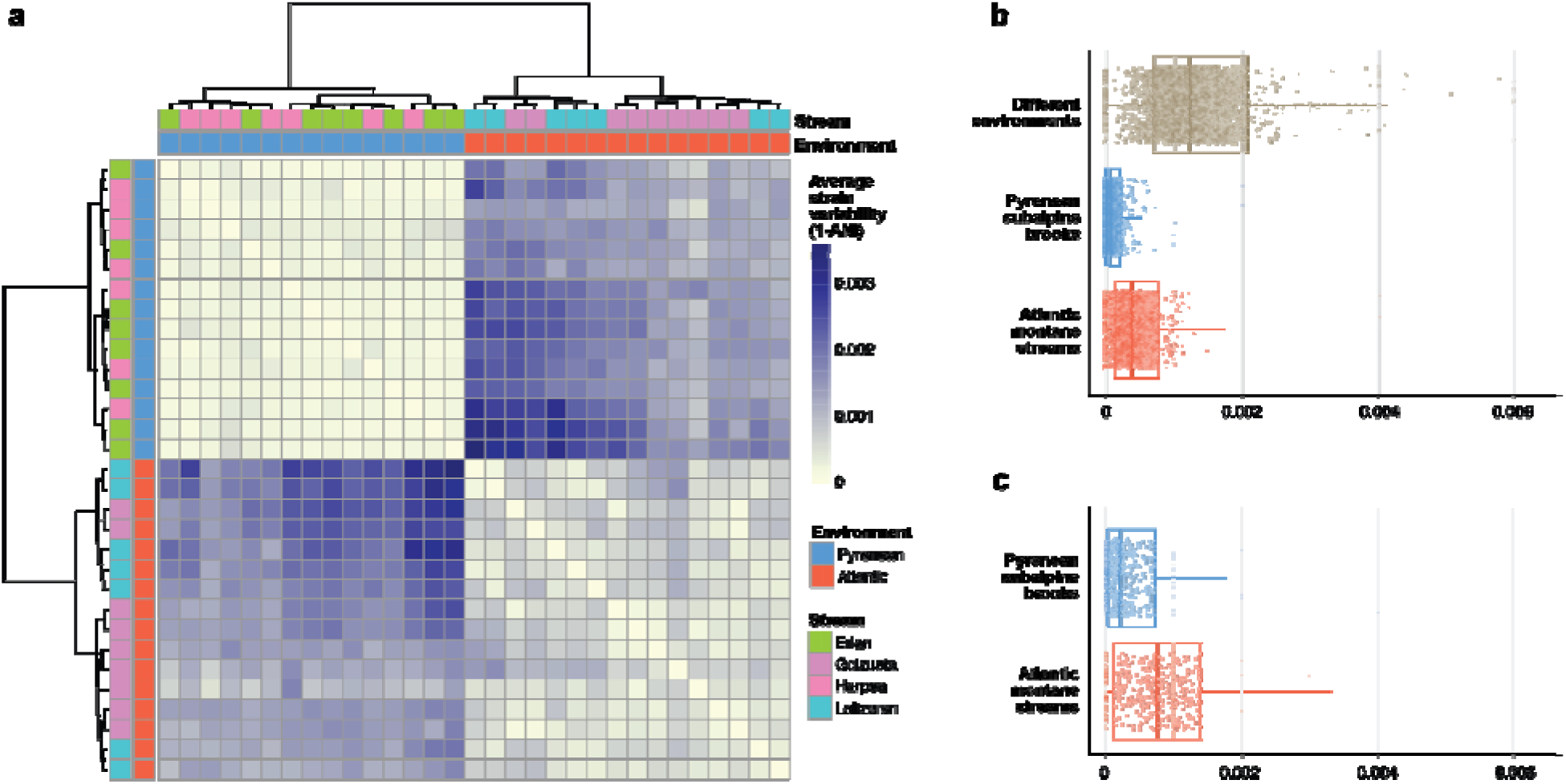
Genome microdiversity. **a)** Heatmap of average pairwise microdiversity (1 – average nucleotide identity) among core genomes across all samples, sorted by hierarchical clustering. Darker tiles indicate larger variability between microbial genomes across samples. **b)** Average pairwise microdiversity of core genomes between samples from different environments, among samples from Pyrenean subalpine brooks, and among samples from Atlantic montane streams. **c)** Average pairwise microdiversity of endemic genomes, comparing Pyrenean versus Atlantic samples.

## Discussion

This first comprehensive characterisation of the gut and skin microbiota of the Pyrenean brook salamander (*Calotriton asper*) unveiled unique insights into the structure and source of functional microbiome variation across populations and environments.

### Uncovering a concealed diversity

To date, the majority of studies profiling amphibian gut and skin microbiotas have relied on 16S amplicon sequencing [72–76]. While this approach has provided an initial glimpse into microbiota composition and structure [77], it is limited by existing sequence diversity, as it relies on previously sequenced and taxonomically annotated 16S rRNA gene references [78]. In contrast, our results highlight how genome-resolved shotgun metagenomics enables the discovery of previously uncharacterised taxa and their corresponding functional capacities, providing a more comprehensive view of host–microbiota relationships and advancing our understanding of microbial ecology in wildlife. Using this approach, we successfully reconstructed 582 bacterial genomes spanning 13 phyla, over 90% of which exhibited an average nucleotide identity below 95% relative to any genome indexed in current databases. These results indicate that the vast majority of bacterial species hosted by *C. asper* are novel to science [79, 80], underscoring how much remains to be discovered about the bacterial diversity of newts and salamanders.

The vast majority (539) of reconstructed bacterial genomes diversity derived from faecal samples, with only 43 genomes reconstructed from skin swab samples. While our domain-adjusted mapping rate (DAMR) analyses indicated near-complete reconstruction of the bacterial genomes present in faecal samples, the same could not be achieved for skin samples. We estimated that nearly half of the bacterial genomic information in the skin samples was likely not represented in our MAG catalogue. This limitation arises because skin swabs typically contain large proportions of host DNA, often exceeding 90% of all reads, together with a diverse array of low-abundance environmental bacteria. Both factors substantially increase sequencing requirements and complicate metagenomic assemblies [81]. The challenge of recovering skin microbiomes is further compounded by the significantly lower amount of DNA present in skin samples compared to faecal samples, which increases the complexity of sequencing library preparation and determines the quality of sequencing [82]. In any case, the pronounced disparity in DAMR values between faecal and skin samples highlights the critical importance of assessing genome recovery success in genome-resolved metagenomics, ensuring the accurate interpretation of results [60].

Despite these caveats, the recovered bacterial communities were markedly distinct. While we cannot entirely rule out the reconstruction of some environmental bacteria, the bacterial communities derived from faecal samples clearly resembled a typical vertebrate gut microbiome, characterised by a high diversity of anaerobic bacteria [83]. This pattern indicates that our careful sampling procedures were effective in capturing the gut-associated microbial community while minimising the contribution of environmental bacteria. The gut microbiome of *C. asper* was dominated by Bacteroidota, Bacillota A, and Pseudomonadota. These phyla were also identified as the most broadly represented in the North American eastern newt (*Notophthalmus viridescens)* [84], suggesting that they may represent a coarse-scale gut microbiota typical of urodeles. The skin microbiome differed markedly from the gut microbiome, displaying significantly lower diversity and an overwhelming dominance of Pseudomonadota. Despite the aforementioned limitations in reconstructing bacterial genomes from skin swabs, the recovered community—dominated by Gammaproteobacteria, Bacteroidia, and Alphaproteobacteria—closely resembled those reported in other newts [85, 86].

### Different microbiomes at different environments

The distinct microbial community structures observed between animals from Atlantic montane streams and Pyrenean subalpine brooks suggest that habitat differences play a significant role in shaping the gut microbiota of *C. asper*. Gut microbiome differences neither affected overall alpha diversity nor metabolic capacity values between habitat types. However, community rearrangements resulted in significant changes in the metabolic properties of the bacterial communities, with significant enrichment in various functions. Notably, these included biosynthesis of several amino acids (e.g., arginine, lysine, phenylalanine, and tryptophan) and their derivatives, as well as enhanced degradation of lipids, polysaccharides, and nitrogen compounds. We hypothesised that these changes could be indicative of a broader dietary niche in salamanders inhabiting Atlantic montane streams, with an expanded exposure to nutrients that produce more metabolic niches for bacteria [87, 88]. However, our dietary analyses did not reveal any notable difference in the coarse-scale dietary items, which were dominated by arthropods and annelids at both environments, with a significant proportion of Tremellales fungi and traces of fabaceus trees, plausibly derivative from ingesting detritus. Regardless of active predation or passive ingestion, these taxa contain different nutrients that could shape gut microbiomes, but their distribution did not show a consistent bias towards a single stream or environment. Interestingly, skin microbiomes, which are not directly related to nutrition, also displayed the same pattern, suggesting that factors other than diet drive the functional differences between environments.

Considering that microbiome profiles often mirror host genomic differences [91, 92], and given that our Atlantic population has not been included in the genetic analysis of *Calotriton asper*[28], we tested whether kmer-based genome skimming of host reads recovered from skin swab samples revealed significant differences between animals from Atlantic montane streams and Pyrenean subalpine brooks. While the resolution of the employed approach is limited, the absence of a significant signal suggests that the genetic differences between the contrasted populations are not drastic, reducing the likelihood that the observed changes were due to genetic differences.

### The contribution of endemic bacteria to functional variation

To further investigate these functional differences, we traced their origins in the gut microbiomes of Pyrenean and Atlantic *C. asper* by categorising the bacteria into core, endemic, and marginal groups. The majority (389 MAGs) were classified as part of the core microbiome. This suggests that, despite geographical separation, genetic isolation of the host populations, and environmental variation, a substantial portion of the *C. asper* microbiome remains consistently associated with the host. These core bacteria constituted the bulk of the microbiome, typically accounting for 80–90% of sequencing reads, indicating a notably robust core microbiome—often associated with good health [93]. A significant subset of bacteria (97 MAGs) was classified as endemic, being both exclusive to and widespread within individuals from only a single environment, and representing 10–15% of the microbiome. Almost the entire diversity of Fusobacteriota—characterised by a high capacity for amino acid degradation—was restricted to endemic bacteria from Atlantic montane streams, contributing to the functional distinctiveness of microbiomes. Lastly, marginal bacteria (53 MAGs), found in only a few individuals within one population, emphasised the stochastic nature of microbiomes in wild animals. Typically comprising less than 5% of the microbiome, these bacteria reflect individual-specific variation not representative of the broader population, though additional sampling is required to confirm their low prevalence across the host population

Partitioning the microbiota also revealed significant differences in the functional properties of the genomes associated with each fraction. We hypothesised that the core bacterial fraction would exhibit smaller average genome sizes, based on the assumption that it comprises taxa closely associated with the host and subject to substantial (pseudo)vertical transmission. Such transmission would be expected to favour bacteria with reduced genomes and limited metabolic capabilities, relying on interactions with other taxa for survival [94]. Contrary to our expectations, the average genome size of core bacteria was not smaller, but in fact significantly larger than that of endemic and marginal taxa. These findings point to an alternative scenario: bacteria with reduced genome size and high metabolic dependency may experience less extensive (pseudo)vertical transmission, as their successful colonisation likely depends on the prior establishment of a structured microbial community capable of providing metabolic by-products essential for their survival [95].

These results also challenge our prior assumption that marginal bacteria are merely opportunistic colonisers, entering gut microbial communities through sporadic events. The similarity in functional profiles between marginal and endemic bacteria suggests that both fractions are largely composed of taxa that integrate into an already complex microbiome, primarily structured by core taxa. These core taxa appear to generate certain metabolic niches more frequently than others, with endemic bacteria occupying the more persistent niches and marginal bacteria associated with the rarer ones [9]. As a result, the ecological significance of marginal bacteria may be greater than previously assumed.

### Increased metabolic capacity and microdiversity at montane streams compared to subalpine brooks

The functional and microdiversity analyses comparing animals across both environments revealed similar patterns. The average metabolic capacity of endemic and marginal taxa from Atlantic animals was higher than that of their Pyrenean counterparts, reflecting the broader observation that skin microbes from Atlantic animals exhibited significantly greater overall metabolic capacities. Additionally, gut bacteria from Atlantic montane stream salamanders displayed higher intraspecific variability than those at Pyrenean subalpine brooks.

While bacterial community diversity analyses are widespread, to our knowledge, no studies have yet characterised community-level variation in genome sizes, metabolic capacities, and microdiversity across environments. Traditional diversity studies in freshwater systems have produced highly context-dependent results, with diversity found to correlate both positively and negatively with elevation [96], largely influenced by regional habitat complexity [97]. Although logistical constraints prevented us from sampling water, detritus, and dietary animal sources—limiting direct comparisons between environmental and gut microbiomes—the consistent patterns observed in both skin and gut bacteria suggest the existence of an environmental microbial pool at Atlantic montane streams with enhanced metabolic capacity and greater strain-level variability compared to Pyrenean subalpine brooks.

These patterns may reflect the higher energy availability, environmental complexity, and overall biodiversity typical of low-elevation streams, which could promote bacterial diversification and functional enrichment [98]. This complexity, in turn, may be mirrored in the microbiomes of newts inhabiting these environments. In our study area, Atlantic montane streams located at lower elevation were characterised by a more structurally complex landscape, comprising a heterogeneous mosaic of forests and small agricultural plots. In contrast, Pyrenean subalpine brooks were situated in more uniform open meadows.

### Potential impact of functional differences on salamander biology

Ultimately, variation in the presumed availability and recruitment of bacteria with diverse functional capacities across environments may influence salamander biology—potentially enhancing nutrient extraction and energy acquisition from different dietary sources. Gut microbiomes of salamanders from Pyrenean subalpine brooks were enriched for B7, K1, and K2 vitamin biosynthesis pathways, as previously observed in reptiles and mammals at high elevations [99, 100]. The higher vitamin biosynthesis capacity may help offset dietary vitamin deficiencies in a nutritionally poorer environment and potentially contribute to the larger body sizes observed at high-elevation sites [101, 102]. Although animals from Pyrenean subalpine brooks exhibited significantly greater snout–vent lengths, no significant differences in weight were observed, suggesting increased fat accumulation in Atlantic individuals inhabiting lower elevations. This could be supported by the two microbial functions most enriched in Atlantic animals—triglyceride and dicarboxylic acid degradation—both associated with lipid metabolism. While the observed functional differences in the microbiome correspond with phenotypic variation, experimental manipulation will be necessary to determine whether microbial properties causally influence salamander physiology.

## Conclusion

Our study provides the first comprehensive catalogue of bacterial genomes associated with wild salamanders and reveals strong ecological patterns through analyses of microbiota partitioning, metabolic capacity, and microdiversity–patterns that were not captured by traditional diversity metrics or taxonomic enrichment analyses. Our results demonstrated that the functional differences across environments were largely driven by endemic bacteria recruited locally. In addition, both functional capacities and intraspecific microdiversity of bacteria differed between contrasting environments, with salamanders from low-elevation Atlantic streams harbouring more diverse and functionally richer microbiomes than their Pyrenean counterparts. The consistency of these patterns in both gut and skin microbiomes suggests that environment-specific factors shape the pool of available bacterial functions, raising the question of whether and how salamanders selectively filter this pool to assemble gut and skin microbiomes that are at least neutral—or potentially beneficial—to their fitness.

Our work, conducted as part of the Earth Hologenome Initiative [27], demonstrates the feasibility of applying genome-resolved metagenomic analyses to wild vertebrates, including the herpetofauna [81]. Amphibians and reptiles have proven well-suited for such investigations, which can yield deep insights into the functional roles of host-associated microorganisms.

Ultimately, this knowledge may prove instrumental in understanding the conservation status of amphibian species and improving captive breeding and translocation efforts aimed at reversing global declines.

## Data accessibility

The raw data and microbial genome sequences were published as part of the 1st EHI Data Release [103], under Bioproject PRJEB76898. The full code that enables reproducing all the analyses included in this manuscript, as well as the accession numbers of the raw data sets, can be found in a dedicated Github repository (https://github.com/alberdilab/calotriton_metagenomics), with a snapshot stored in Zenodo with doi: 10.5281/zenodo.16908317.

## Acknowledgements

This study was funded by the Carlsberg Foundation (grant CF20-0460 awarded to A.A.) and the Danish National Research Foundation (grant DNRF143). We extend our sincere thanks to the Earth Hologenome Initiative (EHI) team for their support, and to the wildlife managers and rangers of Nafarroa and Gipuzkoa, whose collaboration was essential for the successful completion of fieldwork.

