## Supplementary material for "Beyond the core: endemic microbiota drive functional and microdiversity differences across salamander populations"

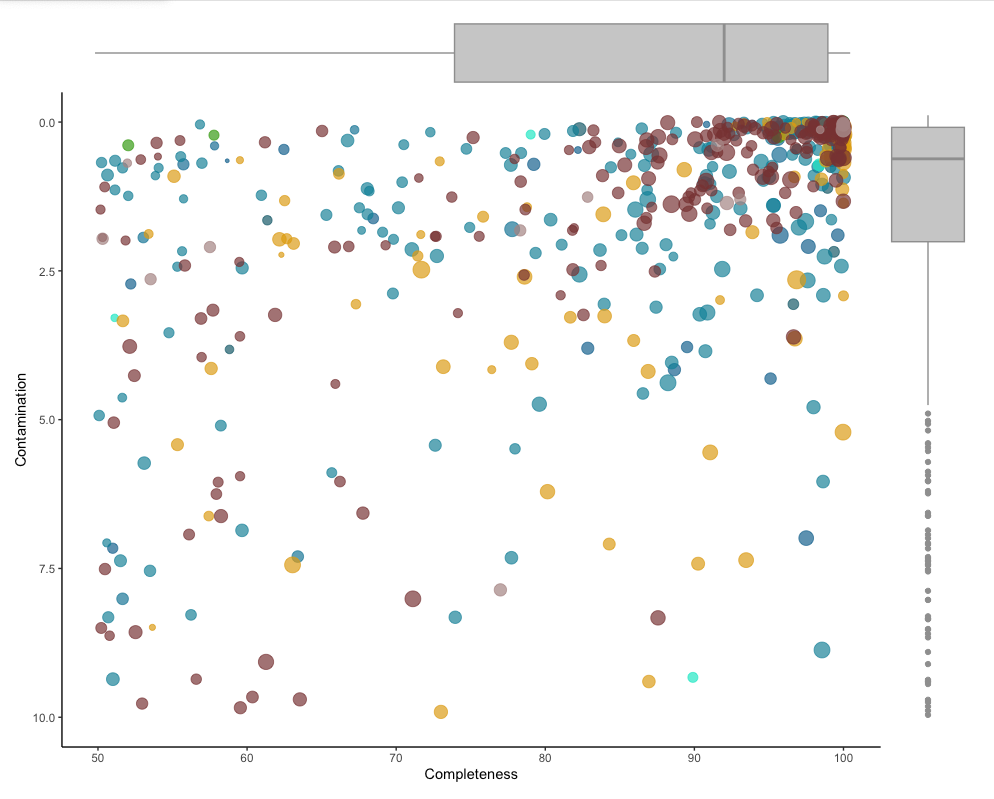

**Figure S1.** Completeness and contamination of genomes reconstructed from faecal samples.

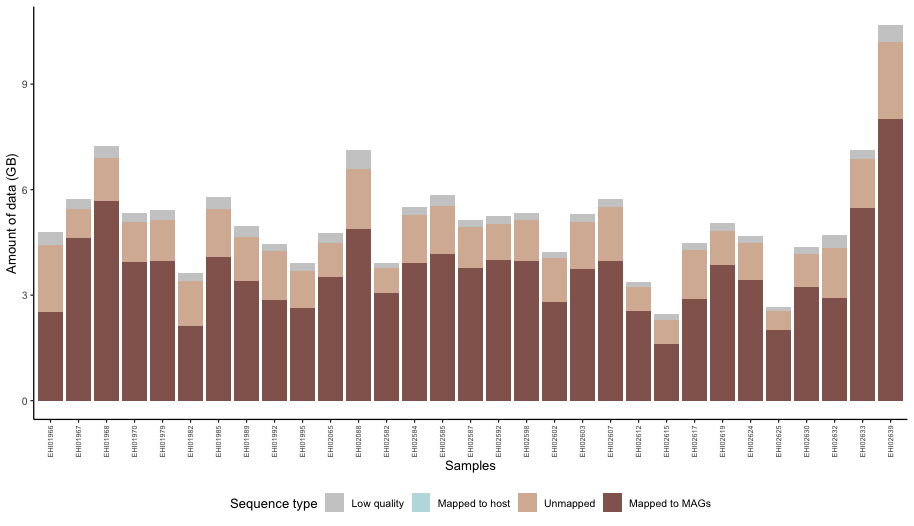

**Figure S2.** Quantitative classification of the sequencing data in faecal samples.

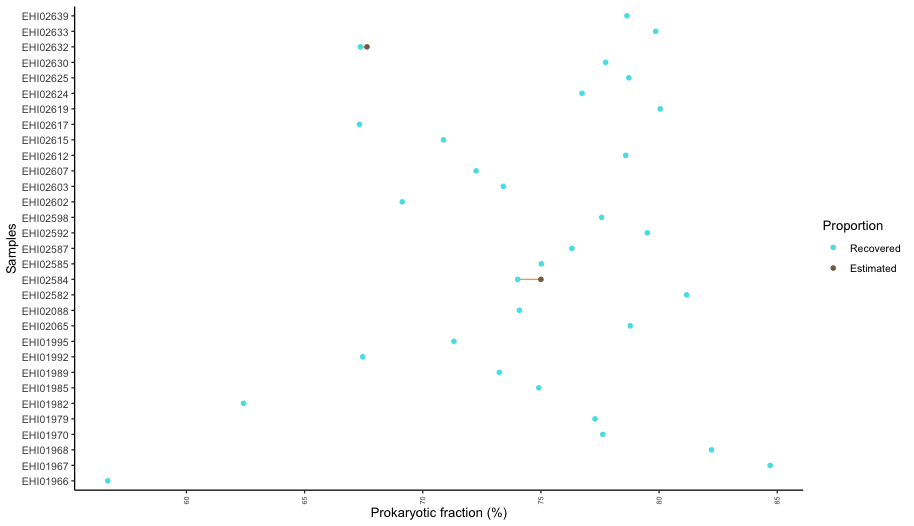

**Figure S3.** Overview of the estimated and recovered fraction of DNA sequences belonging to bacteria and archaea in faecal samples.

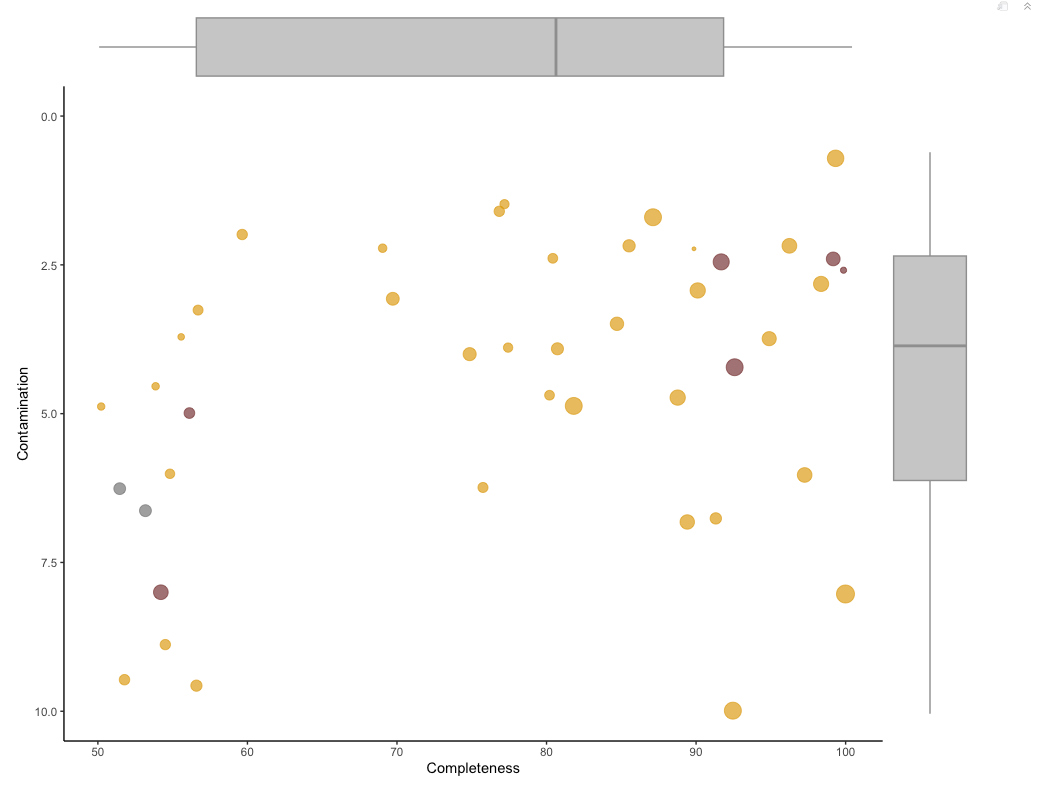

**Table S4.** Completeness and contamination of genomes reconstructed from skin swabs.

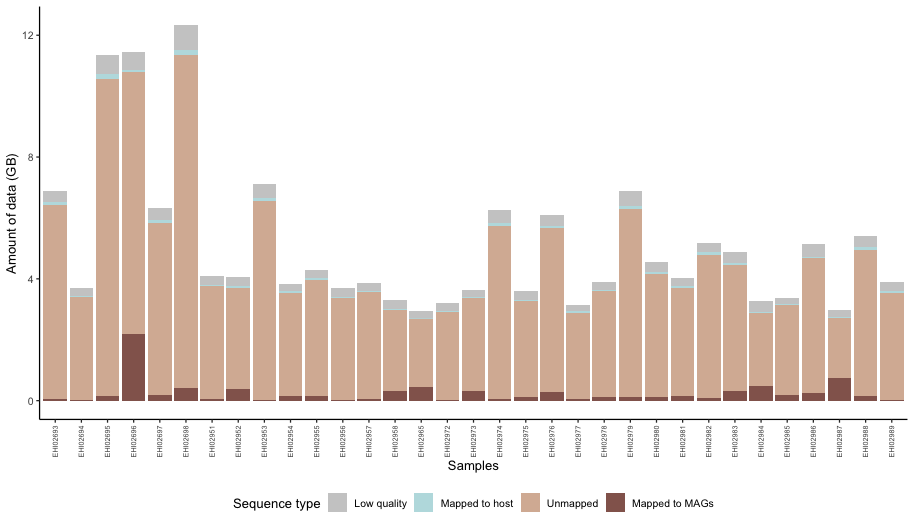

**Figure S5.** Quantitative classification of the sequencing data in skin swabs.

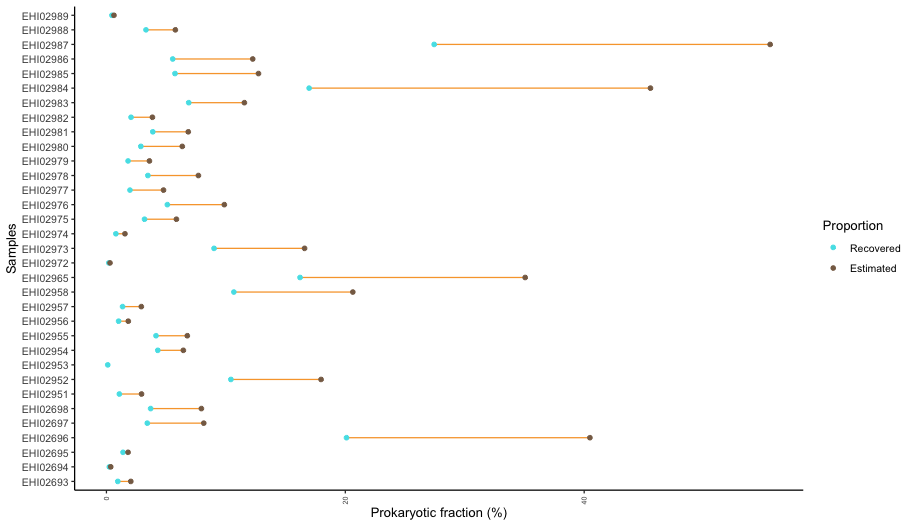

**Figure S6.** Overview of the estimated and recovered fraction of DNA sequences belonging to bacteria and archaea in skin swab samples.

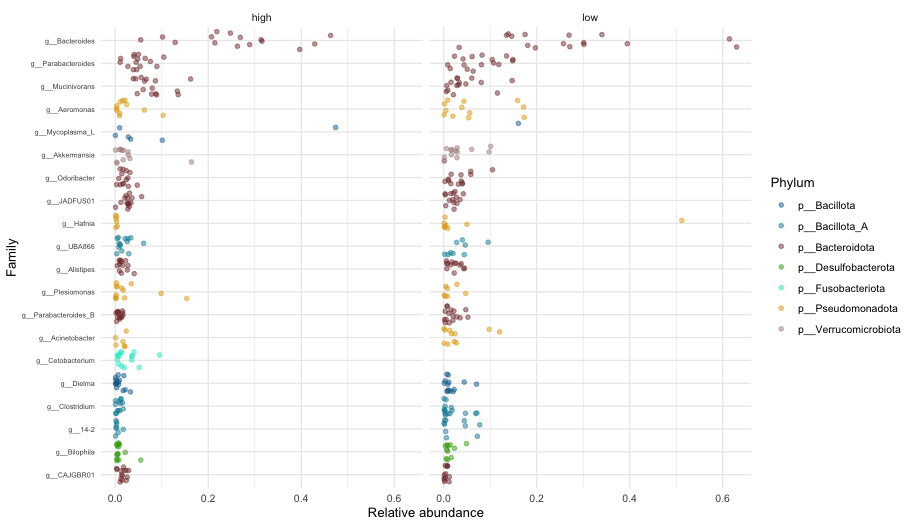

**Figure S7.** Relative abundances of the most abundant gut microbiome genera in animals captured at high and low elevation.

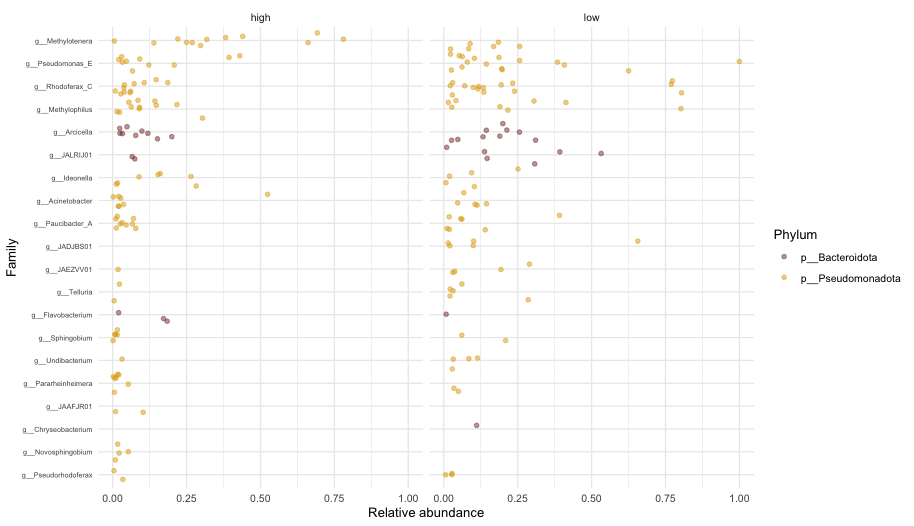

**Figure S8.** Relative abundances of the most abundant skin microbiome genera in animals captured at high and low elevation.

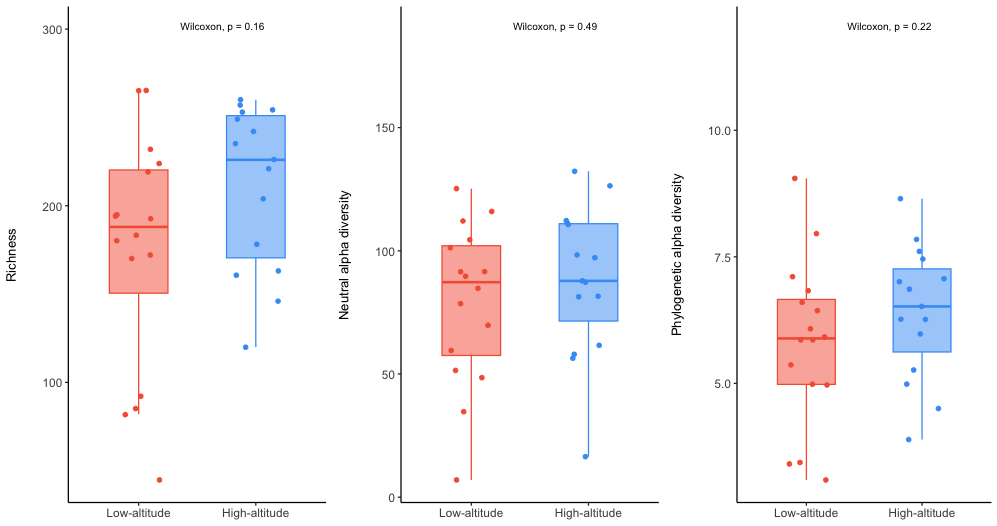

**Figure S9.** Richness, neutral and phylogenetic diversities of gut microbiomes at low and high elevations.

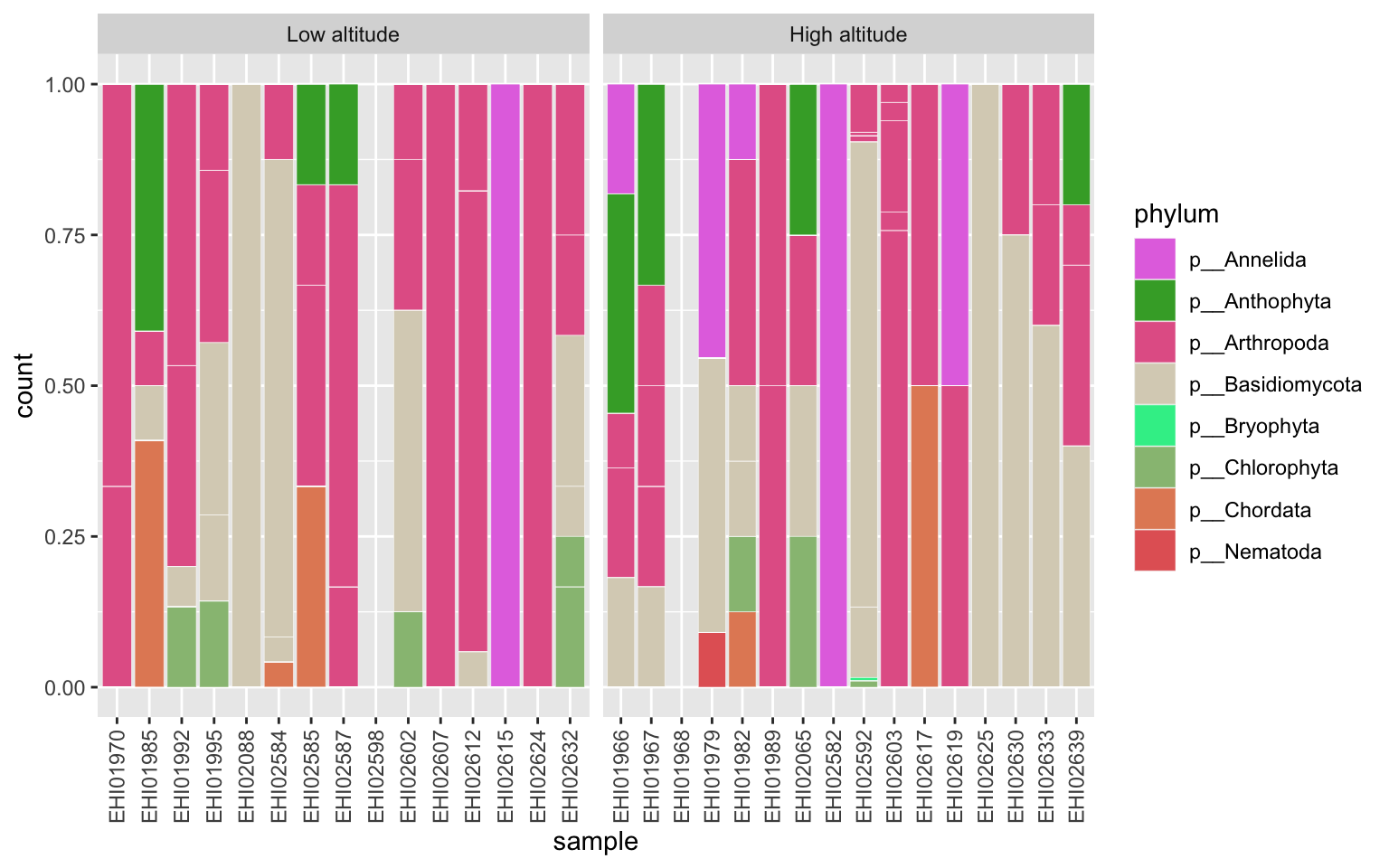

**Figure S10.** Relative read abundance mapped to the marker genes of different orders coloured by phyla.

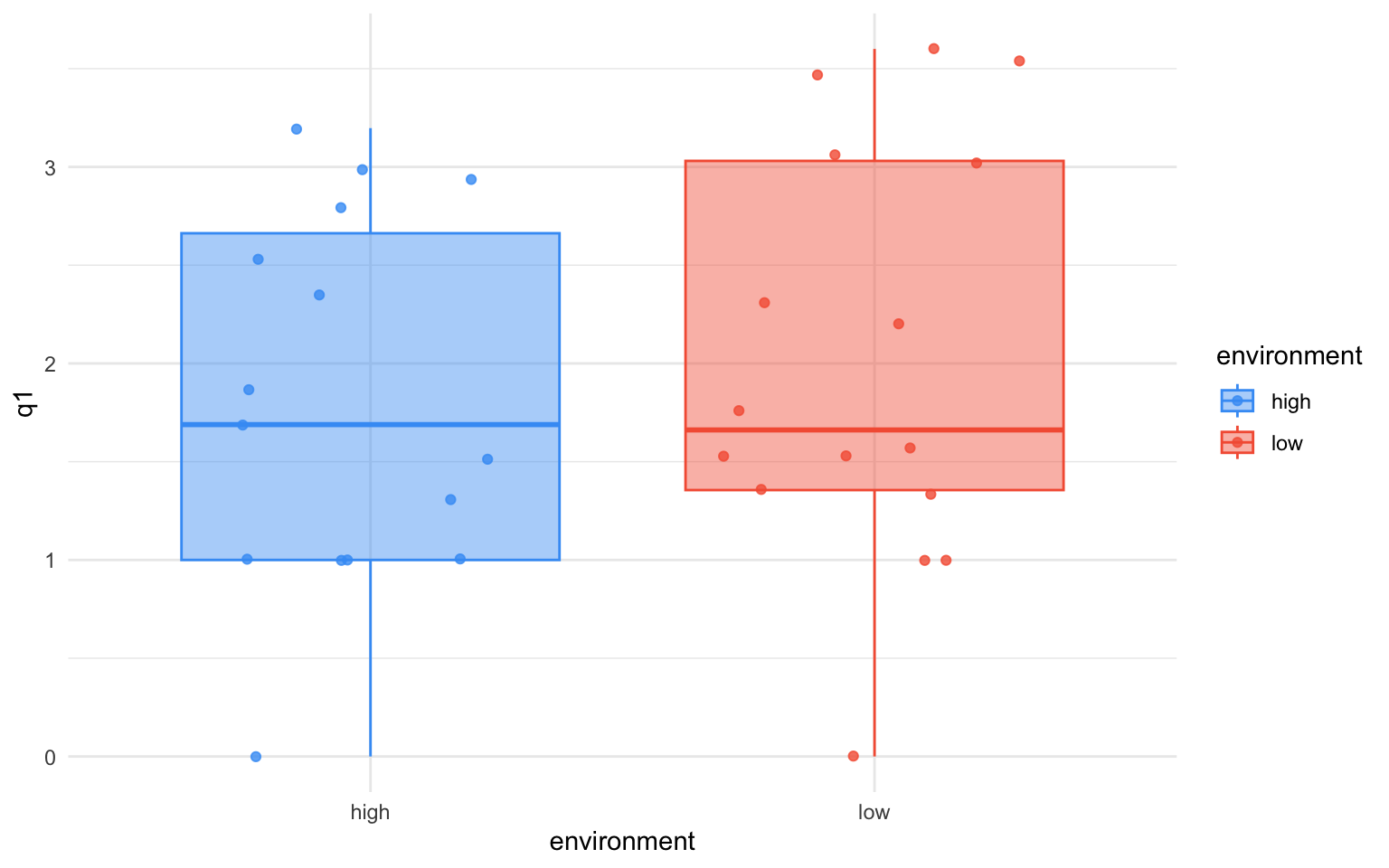

**Figure S11.** Dietary diversity of high and low-elevation animals.

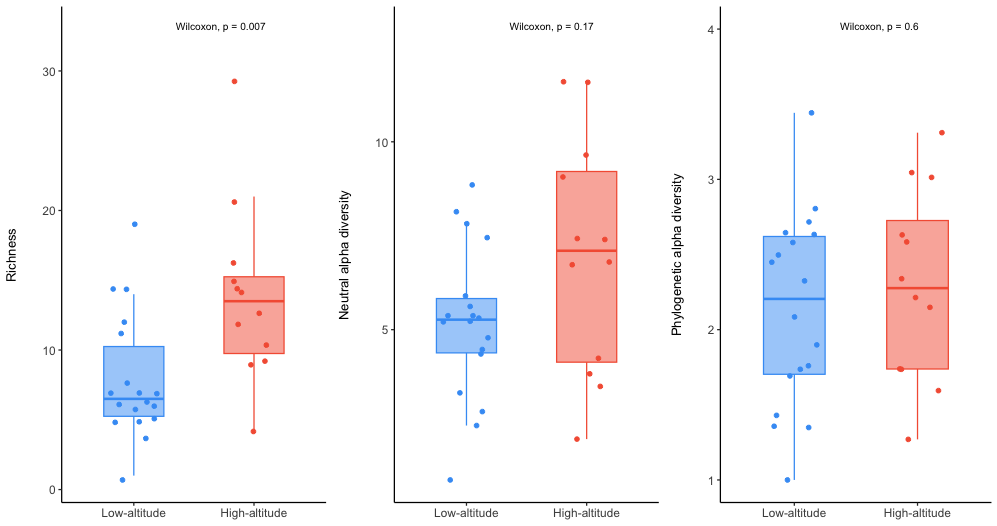
**Figure S12.** Richness, neutral and phylogenetic diversities of skin microbiomes at low and high elevations.

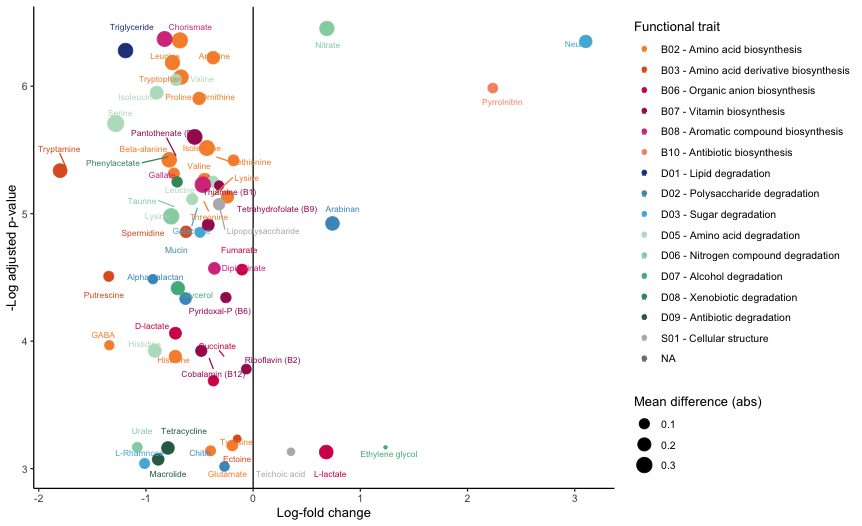

**Figure S13.** Enrichment of functional capacities in low-elevation (left) and high-elevation (right) skin microbiomes.

Table S1. Overview of the genome-inferred functional traits.

| **Code_pathway** | **Code_compound** | **Code_function** | **Type** | **Function** | **Compound** | **Definition** | **module_id** |
| --- | --- | --- | --- | --- | --- | --- | --- |
| B010101 | B0101 | B01 | Biosynthesis | Nucleic acid biosynthesis | Inosinic acid (IMP) | K00764 (K01945,K11787,K11788,K13713) (K00601,K11175,K08289,K11787,K01492) (K01952,(K23269+K23264+K23265),(K23270+K23265)) (K01933,K11787,(K11788 (K01587,K11808,(K01589 K01588)))) (K01923,K01587,K13713) K01756 (K00602,(K01492,(K06863 K11176))) | M00048 |
| B010201 | B0102 | B01 | Biosynthesis | Nucleic acid biosynthesis | Uridylic acid (UMP) | (K11540,((K11541 K01465),((K01954,(K01955+K01956)) ((K00609+K00610),K00608) K01465))) (K00226,K00254,K17828) (K13421,(K00762 K01591)) | M00051 |
| B010301 | B0103 | B01 | Biosynthesis | Nucleic acid biosynthesis | UDP/UTP | K13800,K13809,K09903 | M00052 |
| B010401 | B0104 | B01 | Biosynthesis | Nucleic acid biosynthesis | CDP/CTP | (K00940,K18533) K01937 | M00052 |
| B010501 | B0105 | B01 | Biosynthesis | Nucleic acid biosynthesis | ADP/ATP | K01939 K01756 (K00939,K18532,K18533,K00944) K00940 | M00049 |
| B010601 | B0106 | B01 | Biosynthesis | Nucleic acid biosynthesis | GDP/GTP | K00088 K01951 K00942 (K00940,K18533) | M00050 |
| B020401 | B0204 | B02 | Biosynthesis | Amino acid biosynthesis | Serine | K00058 K00831 (K01079,K02203,K22305,K25528) | M00020 |
| B020501 | B0205 | B02 | Biosynthesis | Amino acid biosynthesis | Threonine | (K00928,K12524,K12525,K12526) K00133 (K00003,K12524,K12525) (K00872,K02204,K02203) K01733 | M00018 |
| B020601 | B0206 | B02 | Biosynthesis | Amino acid biosynthesis | Cysteine | (K00640,K23304) (K01738,K13034,K17069) | M00021 |
| B020602 | B0206 | B02 | Biosynthesis | Amino acid biosynthesis | Cysteine | (K01697,K10150) K01758 | M00338 |
| B020603 | B0206 | B02 | Biosynthesis | Amino acid biosynthesis | Cysteine | K00789 K17462 K01243 K07173 K17216 K17217 | M00609 |
| B020701 | B0207 | B02 | Biosynthesis | Amino acid biosynthesis | Methionine | (K00928,K12524,K12525) K00133 (K00003,K12524,K12525) (K00651,K00641) K01739 (K01760,K14155) (K00548,K24042,K00549) | M00017 |
| B020801 | B0208 | B02 | Biosynthesis | Amino acid biosynthesis | Valine | (K01652+(K01653,K11258)) K00053 K01687 K00826 | M00019 |
| B020901 | B0209 | B02 | Biosynthesis | Amino acid biosynthesis | Isoleucine | (K01703+K01704) K00052 | M00535 |
| B020902 | B0209 | B02 | Biosynthesis | Amino acid biosynthesis | Isoleucine | (K17989,K01754) (K01652+(K01653,K11258)) K00053 K01687 K00826 | M00570 |
| B021001 | B0210 | B02 | Biosynthesis | Amino acid biosynthesis | Leucine | K01649 (K01702,(K01703+K01704)) K00052 | M00432 |
| B021101 | B0211 | B02 | Biosynthesis | Amino acid biosynthesis | Lysine | (K00928,K12524,K12525,K12526) K00133 K01714 K00215 K00674 (K00821,K14267) K01439 K01778 (K01586,K12526) | M00016 |
| B021102 | B0211 | B02 | Biosynthesis | Amino acid biosynthesis | Lysine | K00928 K00133 K01714 K00215 K05822 K00841 K05823 K01778 K01586 | M00525 |
| B021103 | B0211 | B02 | Biosynthesis | Amino acid biosynthesis | Lysine | (K00928,K12524,K12525,K12526) K00133 K01714 K00215 K03340 (K01586,K12526) | M00526 |
| B021104 | B0211 | B02 | Biosynthesis | Amino acid biosynthesis | Lysine | (K00928,K12524,K12525,K12526) K00133 K01714 K00215 K10206 K01778 (K01586,K12526) | M00527 |
| B021105 | B0211 | B02 | Biosynthesis | Amino acid biosynthesis | Lysine | K01655 ((K17450 K01705),(K16792+K16793)) K05824 | M00433 |
| B021106 | B0211 | B02 | Biosynthesis | Amino acid biosynthesis | Lysine | K05827 K05828 K05829 K05830 K05831 | M00031 |
| B021201 | B0212 | B02 | Biosynthesis | Amino acid biosynthesis | Arginine | K00611 K01940 (K01755,K14681) | M00844 |
| B021202 | B0212 | B02 | Biosynthesis | Amino acid biosynthesis | Arginine | K22478 K00145 K00821 K09065 K01438 K01940 K01755 | M00845 |
| B021301 | B0213 | B02 | Biosynthesis | Amino acid biosynthesis | Proline | ((K00931 K00147),K12657) K00286 | M00015 |
| B021401 | B0214 | B02 | Biosynthesis | Amino acid biosynthesis | Glutamate | K00673 K01484 K00840 K06447 K05526 | M00879 |
| B021402 | B0214 | B02 | Biosynthesis | Amino acid biosynthesis | Glutamate | K01745 K01712 K01468 (K01479,K00603,K13990,(K05603 K01458)) | M00045 |
| B021501 | B0215 | B02 | Biosynthesis | Amino acid biosynthesis | Histidine | K00765 ((K01523 K01496),K11755,K14152) (K01814,K24017) ((K02501+K02500),K01663) ((K01693 K00817 (K04486,K05602,K18649)),(K01089 K00817)) (K00013,K14152) | M00026 |
| B021601 | B0216 | B02 | Biosynthesis | Amino acid biosynthesis | Tryptophan | ((((K01657+K01658),K13503,K13501,K01656) K00766),K13497) (((K01817,K24017) (K01656,K01609)),K13498,K13501) ((K01695+(K01696,K06001)),K01694) | M00023 |
| B021701 | B0217 | B02 | Biosynthesis | Amino acid biosynthesis | Phenylalanine | (((K01850,K04092,K14187,K04093,K04516,K06208,K06209) (K01713,K04518,K05359)),K14170) (K00832,K00838) | M00024 |
| B021801 | B0218 | B02 | Biosynthesis | Amino acid biosynthesis | Tyrosine | (((K01850,K04092,K14170,K04093,K04516,K06208,K06209) (K04517,K00211)),K14187) (K00832,K00838) | M00025 |
| B021802 | B0218 | B02 | Biosynthesis | Amino acid biosynthesis | Tyrosine | (K01850,K04092,K14170) (K00832,K15849) (K00220,K24018,K15227) | M00040 |
| B021901 | B0219 | B02 | Biosynthesis | Amino acid biosynthesis | GABA | K09470 K09471 K09472 K09473 | M00136 |
| B022001 | B0220 | B02 | Biosynthesis | Amino acid biosynthesis | Beta-alanine | (K00207,(K17722+K17723)) K01464 (K01431,K06016) | M00046 |
| B022002 | B0220 | B02 | Biosynthesis | Amino acid biosynthesis | Beta-alanine | 6.2.1.17 1.3.8.1 4.2.1.116 3.1.2.4 1.1.1.59 2.6.1.18 | PWY-3941 |
| B022101 | B0221 | B02 | Biosynthesis | Amino acid biosynthesis | Ornithine | (K00618,K00619,K14681,K14682,K00620,K22477,K22478) (((K00930,K22478) K00145),K12659) (K00818,K00821) (K01438,K14677,K00620) | M00028 |
| B022102 | B0221 | B02 | Biosynthesis | Amino acid biosynthesis | Ornithine | K19412 K05828 K05829 K05830 K05831 | M00763 |
| B022103 | B0221 | B02 | Biosynthesis | Amino acid biosynthesis | Ornithine | 2.3.1.1 2.7.2.8 1.2.1.38 2.6.1.11 3.5.1.16 | GLUTORN-PWY |
| B030201 | B0302 | B03 | Biosynthesis | Amino acid derivative biosynthesis | Betaine | 1.1.99.1 1.2.1.8 | BETSYN-PWY |
| B030202 | B0302 | B03 | Biosynthesis | Amino acid derivative biosynthesis | Betaine | 2.1.1.156 2.1.1.157 | P541-PWY |
| B030301 | B0303 | B03 | Biosynthesis | Amino acid derivative biosynthesis | Ectoine | K00928 K00133 K00836 K06718 K06720 | M00033 |
| B030701 | B0307 | B03 | Biosynthesis | Amino acid derivative biosynthesis | Spermidine | (K01583,K01584,K01585,K02626) K01480 | M00133 |
| B030901 | B0309 | B03 | Biosynthesis | Amino acid derivative biosynthesis | Putrescine | K01476 K01581 | M00134 |
| B031001 | B0310 | B03 | Biosynthesis | Amino acid derivative biosynthesis | Tryptamine | 4.1.1.28,4.1.1.105 | PWY-3181 |
| B040101 | B0401 | B04 | Biosynthesis | SCFA biosynthesis | Acetate | (K00625,K13788,K15024) K00925 | M00579 |
| B040103 | B0401 | B04 | Biosynthesis | SCFA biosynthesis | Acetate | K01067 | NA |
| B040104 | B0401 | B04 | Biosynthesis | SCFA biosynthesis | Acetate | 5.4.3.2 5.4.3.3 1.4.1.11 2.3.1.247 1.3.1.109 2.8.3.9 (2.3.1.9,2.3.1.16) 2.3.1.8 (2.7.2.1,2.7.2.15) | P163-PWY |
| B040105 | B0401 | B04 | Biosynthesis | SCFA biosynthesis | Acetate | 1.21.4.2 (2.7.2.1,2.7.2.15) | PWY-8015 |
| B040106 | B0401 | B04 | Biosynthesis | SCFA biosynthesis | Acetate | (1.2.7.1,1.2.1.104) 2.3.1.8 (2.7.2.1,2.7.2.15) | PWY-5100, P41-PWY |
| B040201 | B0402 | B04 | Biosynthesis | SCFA biosynthesis | Butyrate | 1.2.7.1 (2.3.1.9,2.3.1.16) 1.1.1.35 4.2.1.150 1.3.1.109 2.3.1.19 (2.7.2.7,2.7.2.14) | CENTFERM-PWY |
| B040202 | B0402 | B04 | Biosynthesis | SCFA biosynthesis | Butyrate | 2.3.1.8 (2.7.2.1,2.7.2.15) (2.8.3.1,2.8.3.8) | PWY-5676 |
| B040203 | B0402 | B04 | Biosynthesis | SCFA biosynthesis | Butyrate | (2.3.1.9,2.3.1.16) 1.1.1.36 4.2.1.55 1.3.1.109 (2.8.3.1,2.8.3.8) | PWY-5676 |
| B040204 | B0402 | B04 | Biosynthesis | SCFA biosynthesis | Butyrate | 1.4.1.2 1.1.1.399 2.8.3.12 4.2.1.167 7.2.4.5 1.3.1.109 (2.8.3.1,2.8.3.8) | PWY-8190 |
| B040205 | B0402 | B04 | Biosynthesis | SCFA biosynthesis | Butyrate | 5.4.3.2 5.4.3.3 1.4.1.11 2.3.1.247 1.3.1.109 2.8.3.9 | P163-PWY |
| B040206 | B0402 | B04 | Biosynthesis | SCFA biosynthesis | Butyrate | 2.8.3.18 1.2.1.76 1.1.1.61 2.8.3.M6 4.2.1.120 1.3.1.109 (2.8.3.1,2.8.3.8) | PWY-5677 |
| B040207 | B0402 | B04 | Biosynthesis | SCFA biosynthesis | Butyrate | 1.4.1.2 1.1.1.399 2.8.3.12 4.2.1.167 7.2.4.5 1.3.1.109 (2.8.3.1,2.8.3.8) | P162-PWY |
| B040208 | B0402 | B04 | Biosynthesis | SCFA biosynthesis | Butyrate | 1.4.1.2 1.1.1.399 2.8.3.12 4.2.1.167 7.2.4.5 1.3.1.109 (2.8.3.1,2.8.3.8) | PWY-8190 |
| B040301 | B0403 | B04 | Biosynthesis | SCFA biosynthesis | Propionate | (4.2.1.28,1.1.1.1) 1.2.1.87 2.3.1.222 (2.7.2.1,2.7.2.7,2.7.2.14,2.7.2.15) | PWY-7013 |
| B040302 | B0403 | B04 | Biosynthesis | SCFA biosynthesis | Propionate | 4.3.1.19 2.3.1.222 (2.7.2.1,2.7.2.7,2.7.2.14,2.7.2.15) | PWY-5437 |
| B040304 | B0403 | B04 | Biosynthesis | SCFA biosynthesis | Propionate | 2.1.3.1 1.1.1.37 4.2.1.2 1.3.5.1 2.8.3.27 | P108-PWY, PWY-5088 |
| B040305 | B0403 | B04 | Biosynthesis | SCFA biosynthesis | Propionate | 2.8.3.1 4.2.1.54 1.3.1.95 2.8.3.1 | PWY-5494 |
| B040306 | B0403 | B04 | Biosynthesis | SCFA biosynthesis | Propionate | 2.6.1.2 1.1.1.28 2.8.3.1 4.2.1.54 1.3.1.95 2.8.3.1 | PWY-8188 |
| B060101 | B0601 | B06 | Biosynthesis | Organic anion biosynthesis | Succinate | (K01647,K05942) (K01681,K01682) (K00031,K00030) ((((K00164+K00658),K01616)+K00382),(K00174+K00175)) ((K01902+K01903),(K01899+K01900),K18118) ((K00234+K00235+K00236+(K00237,K25801)),(K00239+K00240+K00241),(K00244+K00245+K00246)) (K01676,K01679,(K01677+K01678)) (K00026,K00025,K00024,K00116) | M00009 |
| B060102 | B0601 | B06 | Biosynthesis | Organic anion biosynthesis | Succinate | ((((K00164+K00658),K01616)+K00382),K00174) ((K01902+K01903),(K01899+K01900),K18118) ((K00234+K00235+K00236+(K00237,K25801)),(K00239+K00240+K00241),(K00244+K00245+K00246)) (K01676,K01679,(K01677+K01678)) (K00026,K00025,K00024,K00116) | M00011 |
| B060103 | B0601 | B06 | Biosynthesis | Organic anion biosynthesis | Succinate | K01580 (K13524,K07250,K00823,K16871) (K00135,K00139,K17761) | M00027 |
| B060104 | B0601 | B06 | Biosynthesis | Organic anion biosynthesis | Succinate | (K00169+K00170+K00171+K00172) K01007 K01595 K00024 (K01677+K01678) (K00239+K00240) (K01902+K01903) (K15038,K15017) K14465 (K14467,K18861) K14534 K15016 K00626 | M00374 |
| B060105 | B0601 | B06 | Biosynthesis | Organic anion biosynthesis | Succinate | (K02160+K01961+K01962+K01963) K14468 K14469 K15052 K05606 (K01847,(K01848+K01849)) (K14471+K14472) (K00239+K00240+K00241) K01679 K08691 K14449 K14470 K09709 | M00376 |
| B060106 | B0601 | B06 | Biosynthesis | Organic anion biosynthesis | Succinate | (K00169+K00170+K00171+K00172) (K01959+K01960) K00024 (K01677+K01678) (K18209+K18210) (K01902+K01903) (K00174+K00175+K00176+K00177) | M00620 |
| B060201 | B0602 | B06 | Biosynthesis | Organic anion biosynthesis | Fumarate | (K01647,K05942) (K01681,K01682) (K00031,K00030) ((((K00164+K00658),K01616)+K00382),(K00174+K00175)) ((K01902+K01903),(K01899+K01900),K18118) ((K00234+K00235+K00236+(K00237,K25801)),(K00239+K00240+K00241),(K00244+K00245+K00246)) (K01676,K01679,(K01677+K01678)) (K00026,K00025,K00024,K00116) | M00009 |
| B060202 | B0602 | B06 | Biosynthesis | Organic anion biosynthesis | Fumarate | ((((K00164+K00658),K01616)+K00382),K00174) ((K01902+K01903),(K01899+K01900),K18118) ((K00234+K00235+K00236+(K00237,K25801)),(K00239+K00240+K00241),(K00244+K00245+K00246)) (K01676,K01679,(K01677+K01678)) (K00026,K00025,K00024,K00116) | M00011 |
| B060203 | B0602 | B06 | Biosynthesis | Organic anion biosynthesis | Fumarate | K01948 K00611 K01940 (K01755,K14681) K01476 | M00029 |
| B060204 | B0602 | B06 | Biosynthesis | Organic anion biosynthesis | Fumarate | (K00815,K00838,K00832,K03334) K00457 K00451 K01800 (K01555,K16171) | M00044 |
| B060205 | B0602 | B06 | Biosynthesis | Organic anion biosynthesis | Fumarate | K00241+(K00242,K18859,K18860)+K00239+K00240 | M00149 |
| B060206 | B0602 | B06 | Biosynthesis | Organic anion biosynthesis | Fumarate | K00244+K00245+K00246+K00247 | M00150 |
| B060207 | B0602 | B06 | Biosynthesis | Organic anion biosynthesis | Fumarate | (K00169+K00170+K00171+K00172) K01007 K01595 K00024 (K01677+K01678) (K00239+K00240) (K01902+K01903) (K15038,K15017) K14465 (K14467,K18861) K14534 K15016 K00626 | M00374 |
| B060208 | B0602 | B06 | Biosynthesis | Organic anion biosynthesis | Fumarate | (K02160+K01961+K01962+K01963) K14468 K14469 K15052 K05606 (K01847,(K01848+K01849)) (K14471+K14472) (K00239+K00240+K00241) K01679 K08691 K14449 K14470 K09709 | M00376 |
| B060209 | B0602 | B06 | Biosynthesis | Organic anion biosynthesis | Fumarate | (K00169+K00170+K00171+K00172) (K01959+K01960) K00024 (K01677+K01678) (K18209+K18210) (K01902+K01903) (K00174+K00175+K00176+K00177) | M00620 |
| B060210 | B0602 | B06 | Biosynthesis | Organic anion biosynthesis | Fumarate | (K18029+K18030) K14974 K18028 K15357 K13995 K01799 | M00622 |
| B060211 | B0602 | B06 | Biosynthesis | Organic anion biosynthesis | Fumarate | K00611 K01940 (K01755,K14681) | M00844 |
| B060212 | B0602 | B06 | Biosynthesis | Organic anion biosynthesis | Fumarate | K22478 K00145 K00821 K09065 K01438 K01940 K01755 | M00845 |
| B060213 | B0602 | B06 | Biosynthesis | Organic anion biosynthesis | Fumarate | 2.6.1.1,2.6.1.5,2.6.1.27,2.6.1.57 1.13.11.27 1.13.11.5 5.2.1.2 3.7.1.2 | TYRFUMCAT-PWY |
| B060301 | B0603 | B06 | Biosynthesis | Organic anion biosynthesis | Citrate | (K01647,K05942) (K01681,K01682) (K00031,K00030) ((((K00164+K00658),K01616)+K00382),(K00174+K00175)) ((K01902+K01903),(K01899+K01900),K18118) ((K00234+K00235+K00236+(K00237,K25801)),(K00239+K00240+K00241),(K00244+K00245+K00246)) (K01676,K01679,(K01677+K01678)) (K00026,K00025,K00024,K00116) | M00009 |
| B060302 | B0603 | B06 | Biosynthesis | Organic anion biosynthesis | Citrate | (K01647,K05942) (K01681,K01682) (K00031,K00030) | M00010 |
| B060303 | B0603 | B06 | Biosynthesis | Organic anion biosynthesis | Citrate | K01647 (K01681,K01682) K01637 (K01638,K19282) (K00026,K00025,K00024) | M00012 |
| B060304 | B0603 | B06 | Biosynthesis | Organic anion biosynthesis | Citrate | K01647 K01681 K00031 K00261 (K19268+K01846) K04835 K19280 K14449 K19281 K19282 K00024 | M00740 |
| B060401 | B0604 | B06 | Biosynthesis | Organic anion biosynthesis | L-lactate | 1.1.1.27 | PWY-5481 |
| B060402 | B0604 | B06 | Biosynthesis | Organic anion biosynthesis | L-lactate | 1.1.1.22 | PWY-6713, PWY0-1317 |
| B060501 | B0605 | B06 | Biosynthesis | Organic anion biosynthesis | D-lactate | 1.1.1.28 | PWY-8274 |
| B070101 | B0701 | B07 | Biosynthesis | Vitamin biosynthesis | Thiamine (B1) | (((K03148+K03154) K03151),(K03150 K03149)) K03147 ((K00941 K00788),K14153,K21219) K00946 | M00127 |
| B070102 | B0701 | B07 | Biosynthesis | Vitamin biosynthesis | Thiamine (B1) | (((K03148+K03154) K03151),(K03153 K03149 K10810)) K03147 K00941 K00788 K00946 | M00895 |
| B070103 | B0701 | B07 | Biosynthesis | Vitamin biosynthesis | Thiamine (B1) | (K22699,K03147) ((K00941 (K00788,K21220)),K21219) K00946 | M00896 |
| B070104 | B0701 | B07 | Biosynthesis | Vitamin biosynthesis | Thiamine (B1) | (K00941 K00788),K14153,K21219 | M00899 |
| B070201 | B0702 | B07 | Biosynthesis | Vitamin biosynthesis | Riboflavin (B2) | (((K01497,K14652) ((K01498 K00082),K11752) (K22912,K20860,K20861,K20862,K21063,K21064)),(K02858,K14652)) K00794 K00793 ((K20884 K22949),K11753) | M00125 |
| B070301 | B0703 | B07 | Biosynthesis | Vitamin biosynthesis | Niacin (B3) | 3.6.1.22 (3.2.2.6,3.2.2.4) 3.4.1.19 | PYRIDNUCSAL-PWY |
| B070401 | B0704 | B07 | Biosynthesis | Vitamin biosynthesis | Pantothenate (B5) | ((K00826 K00606 K00077),K01579) (K01918,K13799) | M00119 |
| B070402 | B0704 | B07 | Biosynthesis | Vitamin biosynthesis | Pantothenate (B5) | ((K00606 K00077),(K13367 K00128)) K01918 | M00913 |
| B070501 | B0705 | B07 | Biosynthesis | Vitamin biosynthesis | Pyridoxal-P (B6) | K03472 K03473 K00831 K00097 K03474 K00275 | M00124 |
| B070502 | B0705 | B07 | Biosynthesis | Vitamin biosynthesis | Pyridoxal-P (B6) | K06215 K08681 | M00916 |
| B070601 | B0706 | B07 | Biosynthesis | Vitamin biosynthesis | Biotin (B7) | K00652 (((K00833,K19563) K01935),K19562) K01012 | M00123 |
| B070602 | B0706 | B07 | Biosynthesis | Vitamin biosynthesis | Biotin (B7) | K00652 K25570 K01935 K01012 | M00950 |
| B070603 | B0706 | B07 | Biosynthesis | Vitamin biosynthesis | Biotin (B7) | K16593 K00652 K19563 K01935 K01012 | M00573 |
| B070604 | B0706 | B07 | Biosynthesis | Vitamin biosynthesis | Biotin (B7) | K01906 K00652 (K00833,K19563) K01935 K01012 | M00577 |
| B070701 | B0707 | B07 | Biosynthesis | Vitamin biosynthesis | Tetrahydrofolate (B9) | (K01495,K09007,K22391) (K01077,K01113,(K08310,K19965)) ((K13939,((K13940,(K01633 K00950)) K00796)),(K01633 K13941)) (K11754,K20457) (K00287,K13998) | M00126 |
| B070702 | B0707 | B07 | Biosynthesis | Vitamin biosynthesis | Tetrahydrofolate (B9) | K14652 K22100 K01633 K13941 K22099 K00287 | M00840 |
| B070801 | B0708 | B07 | Biosynthesis | Vitamin biosynthesis | Cobalamin (B12) | (K02302,((K02303,K13542) (K02304,K24866))) (K02190,K03795,K22011) K03394 (K05934,K13541,K21479) K05936 (K02189,K13541) K02188 K05895 ((K02191 K03399),K00595) K06042 K02224 | M00924 |
| B070802 | B0708 | B07 | Biosynthesis | Vitamin biosynthesis | Cobalamin (B12) | (K02303,K13542) (K03394,K13540) K02229 (K05934,K13540,K13541) K05936 K02228 K05895 K00595 K06042 K02224 K02230+K09882+K09883 | M00925 |
| B070803 | B0708 | B07 | Biosynthesis | Vitamin biosynthesis | Cobalamin (B12) | (K00798,K19221) K02232 (K02225,K02227) K02231 K00768 (K02226,K22316) K02233 | M00122 |
| B070901 | B0709 | B07 | Biosynthesis | Vitamin biosynthesis | Tocopherol/tocotorienol (E) | K09833 (K12502,K18534) K09834 K05928 | M00112 |
| B071001 | B0710 | B07 | Biosynthesis | Vitamin biosynthesis | Phylloquinone (K1) | ((K02552 K02551 K08680 K02549),K14759) (K01911,K14760) K01661 (K19222,K12073) K23094 K17872 K23095 | M00932 |
| B071101 | B0711 | B07 | Biosynthesis | Vitamin biosynthesis | Menaquinone (K2) | K02552 K02551 K08680 K02549 K01911 K01661 K19222 K02548 K03183 | M00116 |
| B071102 | B0711 | B07 | Biosynthesis | Vitamin biosynthesis | Menaquinone (K2) | K11782 K18285 (K18286,K20810) K11783 K11784 K11785 | M00930 |
| B071103 | B0711 | B07 | Biosynthesis | Vitamin biosynthesis | Menaquinone (K2) | K11782 K18285 K18284 K11784 K11785 | M00931 |
| B071201 | B0712 | B07 | Biosynthesis | Vitamin biosynthesis | Ubiquinone (Q10) | (K03181,K18240) K03179 (K03182+K03186) K18800 K00568 K03185 K03183 (K03184,K06134) K00568 | M00117 |
| B071202 | B0712 | B07 | Biosynthesis | Vitamin biosynthesis | Ubiquinone (Q10) | K06125 K06126 K00591 K06127 K06134 K00591 | M00128 |
| B080101 | B0801 | B08 | Biosynthesis | Aromatic compound biosynthesis | Salicylate | 5.4.4.2 4.2.99.21 | PWY-6406 |
| B080201 | B0802 | B08 | Biosynthesis | Aromatic compound biosynthesis | Gallate | 4.2.1.10 | PWY-6707 |
| B080301 | B0803 | B08 | Biosynthesis | Aromatic compound biosynthesis | Chorismate | 4.2.1.10 1.1.1.25 2.7.1.71 2.5.1.19 4.2.3.5 | PWY-6163 |
| B080302 | B0803 | B08 | Biosynthesis | Aromatic compound biosynthesis | Chorismate | 2.5.1.54 3.2.3.4 4.2.1.10 1.1.1.25 2.7.1.71 2.5.1.19 4.2.3.5 | PWY-6163 |
| B080303 | B0803 | B08 | Biosynthesis | Aromatic compound biosynthesis | Chorismate | 2.7.2.4 1.2.1.11 2.2.1.10 1.4.1.24 4.2.1.10 1.1.1.25 2.7.1.71 2.5.1.19 4.2.3.5 | PWY-6165 |
| B080404 | B0804 | B08 | Biosynthesis | Aromatic compound biosynthesis | Dipicolinate | 2.7.2.4 1.2.1.11 4.3.3.7 | PWY-8088 |
| B080501 | B0805 | B08 | Biosynthesis | Aromatic compound biosynthesis | Indole-3-acetate | 1.13.12.3 3.5.1.4 | PWY-3161 |
| B080502 | B0805 | B08 | Biosynthesis | Aromatic compound biosynthesis | Indole-3-acetate | 4.2.1.84 3.5.1.4 | PWY-5025 |
| B080503 | B0805 | B08 | Biosynthesis | Aromatic compound biosynthesis | Indole-3-acetate | 3.5.5.1 | PWY-5026 |
| B080504 | B0805 | B08 | Biosynthesis | Aromatic compound biosynthesis | Indole-3-acetate | (2.6.1.1,2.6.1.27) 4.1.1.74 1.2.3.7 | TRPIAACAT-PWY |
| B080505 | B0805 | B08 | Biosynthesis | Aromatic compound biosynthesis | Indole-3-acetate | (4.1.1.28,4.1.1.105) 1.4.3.4 1.2.3.7 | PWY-3181 |
| B090101 | B0901 | B09 | Biosynthesis | Metallophore biosynthesis | Staphyloferrin | K21898 K23446 K23447 | M00876 |
| B090102 | B0901 | B09 | Biosynthesis | Metallophore biosynthesis | Staphyloferrin | (5.1.1.10,5.1.1.12) 6.3.2.58 6.3.2.57 | PWY-7990 |
| B090103 | B0901 | B09 | Biosynthesis | Metallophore biosynthesis | Staphyloferrin | K23371 K21949 K21721 K23372 K23373 K23374 K23375 | M00875 |
| B090104 | B0901 | B09 | Biosynthesis | Metallophore biosynthesis | Staphyloferrin | 2.7.1.225 2.5.1.140 1.5.1.51 6.3.2.54 4.1.1.117 6.3.2.55 6.3.2.56 | PWY-8008 |
| B090201 | B0902 | B09 | Biosynthesis | Metallophore biosynthesis | Aerobactin | K03897 K03896 K03894 K03895 | M00918 |
| B090202 | B0902 | B09 | Biosynthesis | Metallophore biosynthesis | Aerobactin | 1.14.13.59 2.3.1.102 6.3.2.38 6.3.2.39 | AEROBACTINSYN-PWY |
| B090301 | B0903 | B09 | Biosynthesis | Metallophore biosynthesis | Staphylopine | (5.1.1.10,5.1.1.24) 2.5.1.152 1.5.1.52 | PWY-8007 |
| B100402 | B1004 | B10 | Biosynthesis | Antibiotic biosynthesis | Bacilysin | 5.4.99.5 4.1.1.100 5.3.3.19 ((5.3.3.19 1.3.1.aa),1.3.1.aa) 1.1.1.385 6.3.2.49 | PWY-7626 |
| B100601 | B1006 | B10 | Biosynthesis | Antibiotic biosynthesis | Carbapenem-3-carboxylate | K18317 K18316 K18315 | M00675 |
| B100801 | B1008 | B10 | Biosynthesis | Antibiotic biosynthesis | Clavaminate | K12673 K12674 K12675 K12676 | M00674 |
| B100802 | B1008 | B10 | Biosynthesis | Antibiotic biosynthesis | Clavaminate | 2.5.1.66 6.3.3.4 1.14.11.21 3.5.3.22 1.14.11.21 | PWY-5679 |
| B101101 | B1011 | B10 | Biosynthesis | Antibiotic biosynthesis | Erythromycin | 2.1.1.254 1.14.13.154 | PWY-7108 |
| B101102 | B1011 | B10 | Biosynthesis | Antibiotic biosynthesis | Erythromycin | 2.3.1.94 1.14.15.35 2.4.1.328 2.4.1.278 | PWY-7106 |
| B101202 | B1012 | B10 | Biosynthesis | Antibiotic biosynthesis | Fosfomycin | 5.4.2.9 4.1.1.82 1.1.1.309 2.7.7.104 2.1.1.308 1.11.1.23 | PWY-5757 |
| B101401 | B1014 | B10 | Biosynthesis | Antibiotic biosynthesis | Kanosamine | K18652 K18653 K18654 | M00877 |
| B101402 | B1014 | B10 | Biosynthesis | Antibiotic biosynthesis | Kanosamine | 1.1.1.361 2.6.1.104 3.1.3.92 | PWY8J2-22 |
| B102101 | B1021 | B10 | Biosynthesis | Antibiotic biosynthesis | Novobiocin | 6.3.1.15 2.1.1.284 2.4.1.302 2.1.1.285 2.1.3.12 | PWY-7287 |
| B102201 | B1022 | B10 | Biosynthesis | Antibiotic biosynthesis | Paromamine | 4.2.3.124 2.6.1.100 (1.1.1.329,1.1.99.38) 2.6.1.101 2.4.1.283 3.5.1.112 | PWY-7014,PWY-7022 |
| B102401 | B1024 | B10 | Biosynthesis | Antibiotic biosynthesis | Pentalenolactone | K12250 K15907 K18056 K17747 K18091 K18057 K17476 | M00819 |
| B102402 | B1024 | B10 | Biosynthesis | Antibiotic biosynthesis | Pentalenolactone | 4.2.3.7 1.4.15.32 1.14.11.35 1.1.1.340 1.14.13.170 1.14.11.36 1.14.19.8 | PWY-6915 |
| B102601 | B1026 | B10 | Biosynthesis | Antibiotic biosynthesis | Prodigiosin | ((K21780+K21781) K21782 K21783 K21784 K21785 K21786) (K21428 K21778 K21779) K21787 | M00837 |
| B102801 | B1028 | B10 | Biosynthesis | Antibiotic biosynthesis | Pyocyanin | K13063 K20261 K06998 K20260 K20262 K21103 K20940 | M00835 |
| B102802 | B1028 | B10 | Biosynthesis | Antibiotic biosynthesis | Pyocyanin | 2.1.1.327 1.14.13.218 | PWY-6666 |
| B102901 | B1029 | B10 | Biosynthesis | Antibiotic biosynthesis | Pyrrolnitrin | K14266 K19981 K14257 K19982 | M00790 |
| B104101 | B1041 | B10 | Biosynthesis | Antibiotic biosynthesis | Validamycin A | K19969 K20431 K20432 K20433 K20434 K20435 K20436 K20437 K20438 | M00815 |
| B104102 | B1041 | B10 | Biosynthesis | Antibiotic biosynthesis | Validamycin A | 4.2.3.152 5.1.3.33 2.7.1.214 (2.6.1.M1,2.7.7.91) 2.5.1.135 3.1.3.101 2.4.1.338 1.14.11.52 | PWY-5818 |
| B104201 | B1042 | B10 | Biosynthesis | Antibiotic biosynthesis | Violacein | K20086 (K20087+K20088) K20089 K20090 | M00808 |
| B104202 | B1042 | B10 | Biosynthesis | Antibiotic biosynthesis | Violacein | 1.4.3.23 1.21.98.2 1.14.13.217 1.14.13.224 | PWY-7040 |
| B11101 | B111 | B11 | Biosynthesis | Toxin biosynthesis | Cytolethal distending toxin | K11013 K11014 K11015 | NA |
| B11201 | B112 | B11 | Biosynthesis | Toxin biosynthesis | Vacuolating cytotoxin | K11028 | NA |
| B11301 | B113 | B11 | Biosynthesis | Toxin biosynthesis | Enterotoxin ShET-1 | K11011,K11012 | NA |
| B11401 | B114 | B11 | Biosynthesis | Toxin biosynthesis | Cholera toxin | (K10928 K10929),K10954,K10952 | NA |
| B11501 | B115 | B11 | Biosynthesis | Toxin biosynthesis | Exfoliatin | K11041 | NA |
| B11601 | B116 | B11 | Biosynthesis | Toxin biosynthesis | Insecticidal toxin complex | K11063 K11021 | NA |
| B11701 | B117 | B11 | Biosynthesis | Toxin biosynthesis | Typhoid toxin | K11014 K11023 K19298 | NA |
| B11801 | B118 | B11 | Biosynthesis | Toxin biosynthesis | Pertussis toxin | K11023 K11024 K11025 K11026 K11027 | NA |
| B11901 | B119 | B11 | Biosynthesis | Toxin biosynthesis | Hemolysin | K10948,K11032,K11038,K11016,K11017,K11019,K11039,K11068,K11139 | NA |
| B111001 | B1110 | B11 | Biosynthesis | Toxin biosynthesis | Thiol-activated cytolysin | K11031 | NA |
| B111101 | B1111 | B11 | Biosynthesis | Toxin biosynthesis | RTX toxin | K10953,K11022 | NA |
| B111201 | B1112 | B11 | Biosynthesis | Toxin biosynthesis | Non-hemolytic enterotoxin | K11033,K11034 | NA |
| B111301 | B1113 | B11 | Biosynthesis | Toxin biosynthesis | Hydrogen cyanide | K10814 K10815 K10816 | NA |
| D010101 | D0101 | D01 | Degradation | Lipid degradation | Triglyceride | (K01046,K12298,K16816,K13534,K14073,K14074,K14075,K14076,K22283,K14452,K22284,K14674,K14675,K17900) (K01054,K25824) | M00098 |
| D010102 | D0101 | D01 | Degradation | Lipid degradation | Triglyceride | (3.1.1.3,3.1.1.34) (3.1.1.34,3.1.1.79,3.1.1.116) (3.1.1.23,3.1.1.79) | LIPAS-PWY |
| D010201 | D0102 | D01 | Degradation | Lipid degradation | Fatty acid | (K01897,K15013) (K00232,K00249,K00255,K06445,K09479) (((K01692,K07511,K13767) (K00022,K07516)),K01825,K01782,K07514,K07515,K10527) (K00632,K07508,K07509,K07513) | M00086+M00087 |
| D010301 | D0103 | D01 | Degradation | Lipid degradation | Oleate | 6.2.1.3 1.3.8.8 4.2.1.17 1.1.1.35 2.3.1.16 1.3.8.8 4.2.1.17 1.1.1.35 2.3.1.16 1.3.8.8 4.2.1.17 (1.1.1.35,1.1.1.211) 2.3.1.16 5.3.3.8 4.2.1.74 | PWY0-1337 |
| D010401 | D0104 | D01 | Degradation | Lipid degradation | Dicarboxylic acids | 6.2.1.5 1.3.8.7 4.2.1.17 1.1.1.35 2.3.1.174 | PWY-8354 |
| D020101 | D0201 | D02 | Degradation | Polysaccharide degradation | Cellulose | (3.2.1.4,3.2.1.176,3.2.1.132,3.2.1.73) (3.2.1.176,3.2.1.4,3.2.1.14) (1.14.99.54,1.14.99.56,1.14.99.53) (1.14.99.54,1.14.99.53) 1.14.99.54 (1.14.99.54,1.14.99.56) (1.14.99.54,1.14.99.56,1.14.99.53) (3.2.1.4,3.2.1.8,3.2.1.21,3.2.1.25,3.2.1.45,3.2.1.58,3.2.1.73,3.2.1.74,3.2.1.75,3.2.1.78,3.2.1.91,3.2.1.104,3.2.1.123,3.2.1.132,3.2.1.149,3.2.1.151,3.2.1.164,3.2.1.168,3.2.1.73,3.2.1.39,3.2.1.52,3.2.1.132,3.2.1.146) (3.2.1.4,3.2.1.91) (3.2.1.4,3.2.1.176,3.2.1.132,3.2.1.73) (3.2.1.132,3.2.1.4,3.2.1.73,3.2.1.8,3.2.1.156) (3.2.1.4,3.2.1.6,3.2.1.21,3.2.1.73,3.2.1.74,3.2.1.91,3.2.1.151,3.2.1.165) (3.2.1.8,3.2.1.32,3.2.1.4) 3.2.1.4 (3.2.1.4,3.2.1.151,3.2.1.73,2.4.1.207) (3.2.1.4,3.2.1.151) (3.2.1.4,3.2.1.151,3.2.1.78) (3.2.1.4,3.2.1.8,3.2.1.21,3.2.1.25,3.2.1.45,3.2.1.58,3.2.1.73,3.2.1.74,3.2.1.75,3.2.1.78,3.2.1.91,3.2.1.104,3.2.1.123,3.2.1.132,3.2.1.149,3.2.1.151,3.2.1.164,3.2.1.168,3.2.1.73,3.2.1.39,3.2.1.52,3.2.1.132,3.2.1.146) (3.2.1.4,3.2.1.91) (3.2.1.4,3.2.1.176,3.2.1.132,3.2.1.73) (3.2.1.132,3.2.1.4,3.2.1.73,3.2.1.8,3.2.1.156) (3.2.1.4,3.2.1.6,3.2.1.21,3.2.1.73,3.2.1.74,3.2.1.91,3.2.1.151,3.2.1.165) (3.2.1.8,3.2.1.32,3.2.1.4) 3.2.1.4 (3.2.1.4,3.2.1.151,3.2.1.73,2.4.1.207) (3.2.1.4,3.2.1.151) (3.2.1.4,3.2.1.151,3.2.1.78) | NA |
| D020201 | D0202 | D02 | Degradation | Polysaccharide degradation | Xyloglucan | (3.2.1.4,3.2.1.8,3.2.1.21,3.2.1.25,3.2.1.45,3.2.1.58,3.2.1.73,3.2.1.74,3.2.1.75,3.2.1.78,3.2.1.91,3.2.1.104,3.2.1.123,3.2.1.132,3.2.1.149,3.2.1.151,3.2.1.164,3.2.1.168,3.2.1.73,3.2.1.39,3.2.1.52,3.2.1.132,3.2.1.146) (3.2.1.4,3.2.1.6,3.2.1.21,3.2.1.73,3.2.1.74,3.2.1.91,3.2.1.151,3.2.1.165) (3.2.1.4,3.2.1.151,3.2.1.73,2.4.1.207) (2.4.1.207,3.2.1.103,3.2.1.39,3.2.1.6,3.2.1.73,3.2.1.81,3.2.1.83,3.2.1.151,3.2.1.181,3.2.1.178,3.2.1.35,3.2.1.181) (3.2.1.4,3.2.1.151) (3.2.1.4,3.2.1.151,3.2.1.78) (3.2.1.176,3.2.1.4,3.2.1.14) (3.2.1.4,3.2.1.150,3.2.1.151) (1.14.99.54,1.14.99.56) (3.2.1.20,3.2.1.22,3.2.1.24,3.2.1.84,3.2.1.48,3.2.1.10,3.2.1.177,4.2.2.13,2.4.1.161) (3.2.1.37,3.2.1.55,3.2.1.8,3.2.1.99,3.2.1.145,3.2.1.146) (3.2.1.8,3.2.1.32,3.2.1.4) (3.2.1.23,3.2.1.25,3.2.1.31,3.2.1.55,3.2.1.152,3.2.1.165,3.2.1.37,3.2.1.146) | NA |
| D020301 | D0203 | D02 | Degradation | Polysaccharide degradation | Starch | (3.2.1.1,3.2.1.41,2.4.1.19,3.2.1.54,3.2.1.93,3.2.1.10,3.2.1.133,3.2.1.135,3.2.1.20,3.2.1.60,3.2.1.68,3.2.1.70,3.2.1.98,3.2.1.116,2.4.1.18,5.4.99.16,2.4.1.25,2.4.1.4,2.4.1.7,3.2.1.141,5.4.99.11,5.4.99.15,3.2.1.33,2.4.99.16) 3.2.1.2 (3.2.1.1,3.2.1.22,3.2.1.41,3.2.1.54,2.4.1.18,2.4.1.25) 3.2.1.1 3.2.1.33 (3.2.1.3,3.2.1.70,3.2.1.28,2.4.1.2) (3.2.1.3,3.2.1.20,3.2.1.22) | NA |
| D020401 | D0204 | D02 | Degradation | Polysaccharide degradation | Chitin | (3.2.1.14,3.2.1.17,3.2.1.96) (3.2.1.14,3.2.1.17) (3.2.1.17,4.2.2.n1,3.2.1.14) (3.2.1.17,3.2.1.96) (3.2.1.52,3.2.1.140) (3.2.1.21,3.2.1.37,3.2.1.45,3.2.1.52,3.2.1.55,3.2.1.58,3.2.1.74,3.2.1.120,3.2.1.126) (3.2.1.4,3.2.1.8,3.2.1.21,3.2.1.25,3.2.1.45,3.2.1.58,3.2.1.73,3.2.1.74,3.2.1.75,3.2.1.78,3.2.1.91,3.2.1.104,3.2.1.123,3.2.1.132,3.2.1.149,3.2.1.151,3.2.1.164,3.2.1.168,3.2.1.73,3.2.1.39,3.2.1.52,3.2.1.132,3.2.1.146) (3.2.1.52,3.2.1.35,3.2.1.169) (3.2.1.21,3.2.1.37,3.2.1.45,3.2.1.52) (1.14.99.54,1.14.99.56,1.14.99.53) (1.14.99.54,1.14.99.53) (3.1.1.72,3.5.1.41) | NA |
| D020501 | D0205 | D02 | Degradation | Polysaccharide degradation | Pectin | (3.2.1.15,3.2.1.40,3.2.1.67,3.2.1.82,3.2.1.171,3.2.1.173) (4.2.2.2,4.2.2.9,4.2.2.10) (4.2.2.2,4.2.2.9) (4.2.2.2,4.2.2.9) 4.2.2.2 (4.2.2.23,4.2.2.24) 4.2.2.6 4.2.2.24 4.2.2.23 (3.2.1.40,3.2.1.174) 3.2.1.173 3.1.1.11 3.2.1.172 (3.2.1.122,3.2.1.20,3.2.1.22,3.2.1.86,3.2.1.139,3.2.1.67) 3.1.1.72 (3.2.1.23,3.2.1.25,3.2.1.31,3.2.1.55,3.2.1.152,3.2.1.165,3.2.1.37,3.2.1.146) | NA |
| D020601 | D0206 | D02 | Degradation | Polysaccharide degradation | Alpha galactan | 3.2.1.49 (3.2.1.22,3.2.1.49,3.2.1.94,3.2.1.88) 3.2.1.22 (3.2.1.122,3.2.1.20,3.2.1.22,3.2.1.86,3.2.1.139,3.2.1.67) (3.2.1.20,3.2.1.22,3.2.1.24,3.2.1.84,3.2.1.48,3.2.1.10,3.2.1.177,4.2.2.13,2.4.1.161) (3.2.1.22,3.2.1.49,2.4.1.67,2.4.1.82) (3.2.1.3,3.2.1.20,3.2.1.22) | NA |
| D020701 | D0207 | D02 | Degradation | Polysaccharide degradation | Beta-galactan | 3.2.1.89 (3.2.1.23,3.2.1.25,3.2.1.31,3.2.1.55,3.2.1.152,3.2.1.165,3.2.1.37,3.2.1.146) (3.2.1.23,3.2.1.165) 3.2.1.23 (3.2.1.21,3.2.1.23,3.2.1.25,3.2.1.31,3.2.1.37,3.2.1.38,3.2.1.62,3.2.1.74,3.2.1.85,3.2.1.86,3.2.1.105,3.2.1.108,3.2.1.117,3.2.1.118,3.2.1.119,3.2.1.125,3.2.1.147,3.2.1.149,3.2.1.161,3.2.1.175,3.2.1.182) (3.2.1.23,3.2.1.46) | NA |
| D020801 | D0208 | D02 | Degradation | Polysaccharide degradation | Mixed-Linkage glucans | 3.2.1.71 (3.2.1.8,3.2.1.31,3.2.1.37,3.2.1.38,3.2.1.45,3.2.1.75,3.2.1.136) (3.2.1.39,3.2.1.58,3.2.1.73,3.2.1.175) (3.2.1.4,3.2.1.176,3.2.1.132,3.2.1.73) (3.2.1.132,3.2.1.4,3.2.1.73,3.2.1.8,3.2.1.156) (3.2.1.4,3.2.1.6,3.2.1.21,3.2.1.73,3.2.1.74,3.2.1.91,3.2.1.151,3.2.1.165) (3.2.1.4,3.2.1.8,3.2.1.21,3.2.1.25,3.2.1.45,3.2.1.58,3.2.1.73,3.2.1.74,3.2.1.75,3.2.1.78,3.2.1.91,3.2.1.104,3.2.1.123,3.2.1.132,3.2.1.149,3.2.1.151,3.2.1.164,3.2.1.168,3.2.1.73,3.2.1.39,3.2.1.52,3.2.1.132,3.2.1.146) (3.2.1.58,3.2.1.39) 3.2.1.39 (3.2.1.21,3.2.1.37,3.2.1.45,3.2.1.52,3.2.1.55,3.2.1.58,3.2.1.74,3.2.1.120,3.2.1.126) (2.4.1.207,3.2.1.103,3.2.1.39,3.2.1.6,3.2.1.73,3.2.1.81,3.2.1.83,3.2.1.151,3.2.1.181,3.2.1.178,3.2.1.35,3.2.1.181) (3.2.1.39,3.2.1.58,3.2.1.73,3.2.1.175) (3.2.1.58,3.2.1.39) (3.2.1.21,3.2.1.23,3.2.1.25,3.2.1.31,3.2.1.37,3.2.1.38,3.2.1.62,3.2.1.74,3.2.1.85,3.2.1.86,3.2.1.105,3.2.1.108,3.2.1.117,3.2.1.118,3.2.1.119,3.2.1.125,3.2.1.147,3.2.1.149,3.2.1.161,3.2.1.175,3.2.1.182) | NA |
| D020901 | D0209 | D02 | Degradation | Polysaccharide degradation | Xylans | (3.2.1.8,3.2.1.32,3.2.1.4) (3.2.1.78,3.2.1.100,3.2.1.32,3.2.1.73) (3.2.1.132,3.2.1.4,3.2.1.73,3.2.1.8,3.2.1.156) (3.2.1.4,3.2.1.8,3.2.1.21,3.2.1.25,3.2.1.45,3.2.1.58,3.2.1.73,3.2.1.74,3.2.1.75,3.2.1.78,3.2.1.91,3.2.1.104,3.2.1.123,3.2.1.132,3.2.1.149,3.2.1.151,3.2.1.164,3.2.1.168,3.2.1.73,3.2.1.39,3.2.1.52,3.2.1.132,3.2.1.146) (3.2.1.8,3.2.1.32) (3.2.1.8,3.2.1.31,3.2.1.37,3.2.1.38,3.2.1.45,3.2.1.75,3.2.1.136) (3.2.1.102,3.2.1.8,3.2.1.8) (3.2.1.51,3.2.1.8) (3.2.1.76,3.2.1.37) (3.2.1.21,3.2.1.37,3.2.1.45,3.2.1.52,3.2.1.55,3.2.1.58,3.2.1.74,3.2.1.120,3.2.1.126) (3.2.1.20,3.2.1.22,3.2.1.24,3.2.1.84,3.2.1.48,3.2.1.10,3.2.1.177,4.2.2.13,2.4.1.161) 3.2.1.37 (3.2.1.139,3.2.1.131) | NA |
| D021001 | D0210 | D02 | Degradation | Polysaccharide degradation | Beta-mannan | 3.2.1.78 (3.2.1.78,3.2.1.100,3.2.1.32,3.2.1.73) (3.2.1.4,3.2.1.8,3.2.1.21,3.2.1.25,3.2.1.45,3.2.1.58,3.2.1.73,3.2.1.74,3.2.1.75,3.2.1.78,3.2.1.91,3.2.1.104,3.2.1.123,3.2.1.132,3.2.1.149,3.2.1.151,3.2.1.164,3.2.1.168,3.2.1.73,3.2.1.39,3.2.1.52,3.2.1.132,3.2.1.146) (3.2.1.78,3.2.1.100,3.2.1.32,3.2.1.73) (2.4.1.281,2.4.1.319,2.4.1.320) 3.2.1.78 3.1.1.72 (3.2.1.23,3.2.1.25,3.2.1.31,3.2.1.55,3.2.1.152,3.2.1.165,3.2.1.37,3.2.1.146) | NA |
| D021101 | D0211 | D02 | Degradation | Polysaccharide degradation | Alpha-mannan | 3.2.1.130 (3.2.1.101,3.2.1.20) (3.2.1.101,3.2.1.20) (3.2.1.113,3.2.1.24) (3.2.1.24,3.2.1.113,3.2.1.114,3.2.1.170) (3.2.1.106,3.2.1.84,3.2.1.20,3.2.1.170,3.2.1.208) 3.2.1.113 (2.4.1.281,2.4.1.319,2.4.1.320) | NA |
| D021201 | D0212 | D02 | Degradation | Polysaccharide degradation | Arabinan | (3.2.1.37,3.2.1.55,3.2.1.8,3.2.1.99,3.2.1.145,3.2.1.146) (3.2.1.11,3.2.1.57,3.2.1.95) (3.2.1.37,3.2.1.55,3.2.1.8,3.2.1.99,3.2.1.145,3.2.1.146) (3.2.1.4,3.2.1.8,3.2.1.37,3.2.1.55,3.2.1.73) (3.2.1.21,3.2.1.37,3.2.1.45,3.2.1.52,3.2.1.55,3.2.1.58,3.2.1.74,3.2.1.120,3.2.1.126) (3.2.1.55,3.2.1.37) 3.2.1.55 | NA |
| D021301 | D0213 | D02 | Degradation | Polysaccharide degradation | Mucin | 3.2.1.97 (3.2.1.22,3.2.1.49,3.2.1.94,3.2.1.88) 3.2.1.49 (2.4.1.211,2.4.1.247) (3.2.1.20,3.2.1.22,3.2.1.24,3.2.1.84,3.2.1.48,3.2.1.10,3.2.1.177,4.2.2.13,2.4.1.161) (3.2.1.22,3.2.1.49,2.4.1.67,2.4.1.82) | NA |
| D030101 | D0301 | D03 | Degradation | Sugar degradation | Lactose | 3.2.1.85 5.3.1.26 2.7.1.144 4.1.2.40 | LACTOSECAT-PWY |
| D030201 | D0302 | D03 | Degradation | Sugar degradation | Sucrose | 2.7.1.211 3.2.1.48 2.7.1.4 | SUCUTIL-PWY |
| D030302 | D0303 | D03 | Degradation | Sugar degradation | D-Apiose | 1.1.1.420 3.1.1.115 | PWY-8093 |
| D030401 | D0304 | D03 | Degradation | Sugar degradation | D-Arabinose | 5.3.1.3 2.7.1.47 | DARABCAT-PWY |
| D030402 | D0304 | D03 | Degradation | Sugar degradation | D-Arabinose | 5.3.1.3 2.7.1.51 4.1.2.17 1.2.1.21 | DARABCATK12-PWY |
| D030501 | D0305 | D03 | Degradation | Sugar degradation | D-Mannose | 2.7.1.191 5.3.1.8 | MANNCAT-PWY |
| D030502 | D0305 | D03 | Degradation | Sugar degradation | D-Mannose | 2.7.1.7 5.3.1.8 | WY3O-1743 |
| D030601 | D0306 | D03 | Degradation | Sugar degradation | D-Xylose | 5.3.1.5 2.7.1.17 | XYLCAT-PWY |
| D030602 | D0306 | D03 | Degradation | Sugar degradation | D-Xylose | 1.1.1.9 2.7.1.17 | LARABITOLUTIL-PWY |
| D030603 | D0306 | D03 | Degradation | Sugar degradation | D-Xylose | 1.1.1.424 3.1.1.68 4.2.1.82 4.2.1.141 1.2.1.26 | PWY-6760 |
| D030604 | D0306 | D03 | Degradation | Sugar degradation | D-Xylose | 1.1.1.359 3.1.1.110 4.2.1.82 4.1.2.28 1.1.1.26 2.3.3.9 | PWY-7294 |
| D030605 | D0306 | D03 | Degradation | Sugar degradation | D-Xylose | 1.1.1.175 3.1.1.110 4.2.1.82 4.2.1.1.141 1.2.1.26 | PWY-8020 |
| D030701 | D0307 | D03 | Degradation | Sugar degradation | L-Fucose | 5.1.3.29 5.3.1.25 2.7.1.51 4.1.2.17 | FUCCAT-PWY |
| D030702 | D0307 | D03 | Degradation | Sugar degradation | L-Fucose | 5.1.3.29 4.2.1.68 1.1.1.M68 3.7.1.26 | PWY-8318 |
| D030801 | D0308 | D03 | Degradation | Sugar degradation | L-Rhamnose | 5.1.3.32 5.3.1.14 2.7.1.5 4.1.2.19 | RHAMCAT-PWY |
| D030802 | D0308 | D03 | Degradation | Sugar degradation | L-Rhamnose | (1.1.1.173,1.1.1.378) 3.1.1.65 4.2.1.90 4.1.2.53 | PWY-6713 |
| D030803 | D0308 | D03 | Degradation | Sugar degradation | L-Rhamnose | (1.1.1.173,1.1.1.378) 3.1.1.65 4.2.1.90 1.1.1.401 3.7.1.26 | PWY-6714 |
| D030901 | D0309 | D03 | Degradation | Sugar degradation | Galactose | K01785 K00849 K00965 K01784 | M00632 |
| D031001 | D0310 | D03 | Degradation | Sugar degradation | NeuAc | 4.1.3.3 5.1.3.8 | PWY-7581 |
| D031002 | D0310 | D03 | Degradation | Sugar degradation | NeuAc | 4.1.3.3 2.7.1.60 5.1.3.9 | PWY0-1324 |
| D040101 | D0401 | D04 | Degradation | Protein degradation | Albumin | A1A (S8A C13 (S1A (S8B S26B))) | NA |
| D040201 | D0402 | D04 | Degradation | Protein degradation | Actin | S1A (M10A A1A M35 (M9B S1B S1C C1A (M12A C14A A2A C13 M10C))) | NA |
| D040301 | D0403 | D04 | Degradation | Protein degradation | Collagen | C1A M10A (M12B (S1A A1A (C13 M35 (M12A S1D S8A A2A M4 S8B)))) | NA |
| D040401 | D0404 | D04 | Degradation | Protein degradation | Elastin | M10A (S1A (M23A S01C C1A)) | NA |
| D040501 | D0405 | D04 | Degradation | Protein degradation | Glutelin | S26B S8A S9A | NA |
| D040601 | D0406 | D04 | Degradation | Protein degradation | Keratin | S1D M12A S1A | NA |
| D040701 | D0407 | D04 | Degradation | Protein degradation | Tropomyosin | S1B (C2A C1A S1A (M4 C13 M10A (S1D M12A))) | NA |
| D040801 | D0408 | D04 | Degradation | Protein degradation | Troponin | C2A (A2A M35 C14A) | NA |
| D050101 | D0501 | D05 | Degradation | Amino acid degradation | Serine | 4.3.1.17,4.3.1.18 | SERDEG-PWY |
| D050201 | D0502 | D05 | Degradation | Amino acid degradation | Threonine | 4.3.1.19 | PWY-5437 |
| D050301 | D0503 | D05 | Degradation | Amino acid degradation | Cysteine | 4.4.1.1,4.4.1.28 | LCYSDEG-PWY |
| D050302 | D0503 | D05 | Degradation | Amino acid degradation | Cysteine | 2.6.1.3 2.8.1.2 | PWY-5329 |
| D050401 | D0504 | D05 | Degradation | Amino acid degradation | Methionine | 4.4.1.11 | PWY-701 |
| D050501 | D0505 | D05 | Degradation | Amino acid degradation | Valine | 2.6.1.42 1.2.1.25 1.3.8.5 4.2.1.150 3.1.2.4 1.1.1.31 1.2.1.27 | VALDEG-PWY |
| D050502 | D0505 | D05 | Degradation | Amino acid degradation | Valine | 2.6.1.42 4.1.1.72 1.1.1.1 | PWY-5057 |
| D050601 | D0506 | D05 | Degradation | Amino acid degradation | Isoleucine | 2.6.1.42 1.2.1.25 1.3.8.5 4.2.1.150 1.1.1.178 2.3.1.16 | ILEUDEG-PWY |
| D050602 | D0506 | D05 | Degradation | Amino acid degradation | Isoleucine | 2.6.1.42 1.2.7.7 | PWY-8184 |
| D050701 | D0507 | D05 | Degradation | Amino acid degradation | Leucine | K00826 (((K00166+K00167),K11381)+K09699+K00382) (K00253,K00249) (K01968+K01969) (K05607,K13766) K01640 | M00036 |
| D050801 | D0508 | D05 | Degradation | Amino acid degradation | Lysine | 4.1.1.18 2.6.1.82 1.2.1.19 2.6.1.48 1.2.1.20 1.14.11.64 1.1.5.13 | PWY0-461 |
| D050802 | D0508 | D05 | Degradation | Amino acid degradation | Lysine | 1.13.12.2 3.5.1.30 1.6.1.48 1.2.1.20 2.8.3.13 | PWY-5280 |
| D050803 | D0508 | D05 | Degradation | Amino acid degradation | Lysine | 2.6.1.36 1.2.1.31 | PWY-5298 |
| D050804 | D0508 | D05 | Degradation | Amino acid degradation | Lysine | 4.1.1.18 2.6.1.82 1.2.1.19 2.6.1.48 1.2.1.20 2.8.3.13 | PWY-6328 |
| D050805 | D0508 | D05 | Degradation | Amino acid degradation | Lysine | K01582 K09251 K00137 K07250 K00135 K15737 K15736 | M00956 |
| D050806 | D0508 | D05 | Degradation | Amino acid degradation | Lysine | K00468 K01506 (K14268,K07250) K00135 ((K15737 K15736),(K01041 K00252 (K01692,K01825,K01782) (K01825,K01782) K00626)) | M00957 |
| D050901 | D0509 | D05 | Degradation | Amino acid degradation | Arginine | 4.1.1.19 3.5.3.11 | PWY0-823 |
| D050902 | D0509 | D05 | Degradation | Amino acid degradation | Arginine | 4.1.1.19 3.5.3.12 3.5.1.53 | ARGDEG-III-PWY |
| D050903 | D0509 | D05 | Degradation | Amino acid degradation | Arginine | 1.13.12.1 3.5.1.4 3.5.3.7 | ARGDEG-V-PWY |
| D050904 | D0509 | D05 | Degradation | Amino acid degradation | Arginine | K01476 K01581 | M00134 |
| D050905 | D0509 | D05 | Degradation | Amino acid degradation | Arginine | K00613 K00542 K00933 | M00047 |
| D050906 | D0509 | D05 | Degradation | Amino acid degradation | Arginine | (K01583,K01584,K01585,K02626) K01480 K01611 K00797 | M00133 |
| D051001 | D0510 | D05 | Degradation | Amino acid degradation | Proline | 5.1.1.4 1.21.4.1 | PWY-8186 |
| D051101 | D0511 | D05 | Degradation | Amino acid degradation | Glutamate | 1.4.1.2,1.4.1.3 | GLUTAMATE-DEG1-PWY, PWY-5766 |
| D051102 | D0511 | D05 | Degradation | Amino acid degradation | Glutamate | 2.6.1.1 4.3.1.1 | GLUTDEG-PWY |
| D051103 | D0511 | D05 | Degradation | Amino acid degradation | Glutamate | 1.4.1.2 1.1.1.399 2.8.3.12 4.2.1.167 7.2.4.5 | P162-PWY |
| D051104 | D0511 | D05 | Degradation | Amino acid degradation | Glutamate | 5.4.99.1 4.3.1.2 4.2.1.34 4.1.3.22 | PWY-5087, GLUDEG-II-PWY |
| D051105 | D0511 | D05 | Degradation | Amino acid degradation | Glutamate | 4.1.1.15 | PWY0-1305 |
| D051201 | D0512 | D05 | Degradation | Amino acid degradation | Histidine | K01745 K01712 K01468 (K01479,K00603,K13990,(K05603 K01458)) | M00045 |
| D051301 | D0513 | D05 | Degradation | Amino acid degradation | Tryptophan | (K00453,K00463) (K01432,K14263,K07130) K00486 K01556 K00452 K03392 (K10217,K23234) | M00038 |
| D051302 | D0513 | D05 | Degradation | Amino acid degradation | Tryptophan | 4.1.99.1 | TRYPDEG-PWY |
| D051601 | D0516 | D05 | Degradation | Amino acid degradation | Beta-alanine | 2.6.1.18 1.2.1.18 | PWY-1781 |
| D051602 | D0516 | D05 | Degradation | Amino acid degradation | Beta-alanine | 2.6.1.120 1.1.1.298 | PWY-8120 |
| D051701 | D0517 | D05 | Degradation | Amino acid degradation | Ornithine | 4.1.1.17 ((2.6.1.82 1.2.1.19),(6.3.1.11 1.4.3.M3 1.2.1.00 3.5.1.94)) | ORNDEG-PWY, ORNARGDEG-PWY |
| D051801 | D0518 | D05 | Degradation | Amino acid degradation | GABA | 2.6.1.19 (1.2.1.24,1.2.1.16) | PWY-6535,PWY-6536 |
| D051802 | D0518 | D05 | Degradation | Amino acid degradation | GABA | 2.6.1.19 1.1.1.61 2.8.3.M6 4.2.1.120 1.3.1.109 (2.8.3.1,2.8.3.8) | PWY-5022 |
| D060101 | D0601 | D06 | Degradation | Nitrogen compound degradation | Nitrate | ((K00370+K00371+K00374),(K02567+K02568)) ((K00362+K00363),(K03385+K15876)) | M00530 |
| D060102 | D0601 | D06 | Degradation | Nitrogen compound degradation | Nitrate | 1.7.5.1 1.7.2.1 1.7.2.5 1.7.2.4 | DENITRIFICATION-PWY |
| D060103 | D0601 | D06 | Degradation | Nitrogen compound degradation | Nitrate | 1.9.6.1 1.7.2.2 | PWY-5674 |
| D060105 | D0601 | D06 | Degradation | Nitrogen compound degradation | Nitrate | 1.7.7.2 1.7.7.1 | PWY490-3 |
| D060201 | D0602 | D06 | Degradation | Nitrogen compound degradation | Urea | 6.3.4.6 3.5.1.54 | PWY-5703 |
| D060202 | D0602 | D06 | Degradation | Nitrogen compound degradation | Urea | 3.5.1.5 | PWY-5704 |
| D060301 | D0603 | D06 | Degradation | Nitrogen compound degradation | Urate | 1.7.3.3 3.5.2.17 4.1.1.97 | PWY-5691 |
| D060302 | D0603 | D06 | Degradation | Nitrogen compound degradation | Urate | 1.14.13.113 3.5.2.17 4.1.1.97 | PWY-7394 |
| D060401 | D0604 | D06 | Degradation | Nitrogen compound degradation | GlcNAc | 2.7.1.59 3.5.1.25 3.5.99.6 | PWY-6517 |
| D060402 | D0604 | D06 | Degradation | Nitrogen compound degradation | GlcNAc | 3.5.1.25 3.5.99.6 | GLUAMCAT-PWY |
| D060601 | D0606 | D06 | Degradation | Nitrogen compound degradation | Allantoin | 3.5.2.5 3.5.3.9 3.5.3.26 (1.1.1.350,1.1.1.154) 2.1.3.5 | PWY0-41 |
| D060602 | D0606 | D06 | Degradation | Nitrogen compound degradation | Allantoin | 3.5.2.5 3.5.3.4 4.3.2.3 | PWY-5694 |
| D060603 | D0606 | D06 | Degradation | Nitrogen compound degradation | Allantoin | 3.5.2.5 3.5.3.9 3.5.3.26 4.3.2.3 | PWY-5705 |
| D060701 | D0607 | D06 | Degradation | Nitrogen compound degradation | Creatinine | 3.5.4.21 3.5.2.14 3.5.1.59 1.5.3.1 | PWY-4722 |
| D060801 | D0608 | D06 | Degradation | Nitrogen compound degradation | Betaine | 2.1.1.5 1.5.3.10 (1.5.3.24,1.5.3.1) | PWY-3661 |
| D060901 | D0609 | D06 | Degradation | Nitrogen compound degradation | L-carnitine | 1.14.13.239 1.2.1.4 1.1.1.38 | PWY-3641 |
| D061001 | D0610 | D06 | Degradation | Nitrogen compound degradation | Methylamine | 1.4.9.1 | PWY-6967 |
| D061002 | D0610 | D06 | Degradation | Nitrogen compound degradation | Methylamine | 6.3.4.12 2.1.1.21 1.5.99.5 | PWY-6965 |
| D061101 | D0611 | D06 | Degradation | Nitrogen compound degradation | Phenylethylamine | (1.4.3.4,1.4.3.21) 1.2.1.39 | 2PHENDEG-PWY |
| D061201 | D0612 | D06 | Degradation | Nitrogen compound degradation | Hypotaurine | 2.6.1.77 1.2.1.3 | PWY-7387 |
| D061301 | D0613 | D06 | Degradation | Nitrogen compound degradation | Taurine | 2.6.1.77 | PWY-1263 |
| D061303 | D0613 | D06 | Degradation | Nitrogen compound degradation | Taurine | 2.5.1.55 | TAURINEDEG-PWY |
| D070101 | D0701 | D07 | Degradation | Alcohol degradation | 2,3-Butanediol | 1.1.1.4,1.1.1.76 | PWY-6388 |
| D070201 | D0702 | D07 | Degradation | Alcohol degradation | Ethanol | 1.1.1.1 1.2.1.10 | ETOH-ACETYLCOA-ANA-PWY |
| D070202 | D0702 | D07 | Degradation | Alcohol degradation | Ethanol | 1.1.1.1 1.2.1.3 6.2.1.1 | PWY66-21 |
| D070401 | D0704 | D07 | Degradation | Alcohol degradation | Glycerol | 2.7.1.30 1.1.5.3 | PWY0-381 |
| D070402 | D0704 | D07 | Degradation | Alcohol degradation | Glycerol | 1.1.1.6 2.7.1.29 | PWY-6131 |
| D070403 | D0704 | D07 | Degradation | Alcohol degradation | Glycerol | 4.2.1.30 1.1.1.202 | PWY-6130 |
| D070404 | D0704 | D07 | Degradation | Alcohol degradation | Glycerol | 1.1.1.6 2.7.1.121 | GLYCEROLMETAB-PWY |
| D070501 | D0705 | D07 | Degradation | Alcohol degradation | Propylene glycol | 4.2.1.28 (1.1.1.1,(1.2.1.87 2.3.1.222 (2.7.2.1,2.7.2.7,2.7.2.14,2.7.2.15))) | PWY-7013 |
| D070601 | D0706 | D07 | Degradation | Alcohol degradation | Ethylene glycol | 1.1.1.77 1.2.1.21 | PWY0-1280 |
| D070801 | D0708 | D07 | Degradation | Alcohol degradation | Phytol | 1.1.1.1 1.2.1.3 6.2.1.3 1.3.1.38 | PWY66-389 |
| D070901 | D0709 | D07 | Degradation | Alcohol degradation | Polyvinyl alcohol | 1.1.2.6 | PWY-6464 |
| D080101 | D0801 | D08 | Degradation | Xenobiotic degradation | Toluene | (K15760+K15761+K15763+K15764) K00055 K00141 | M00538 |
| D080103 | D0801 | D08 | Degradation | Xenobiotic degradation | Toluene | K07540 (K07543+K07544) K07545 K07546 (K07547+K07548) (K07549+K07550) | M00418 |
| D080201 | D0802 | D08 | Degradation | Xenobiotic degradation | Xylene | (K15757+K15758) K00055 K00141 | M00537 |
| D080402 | D0804 | D08 | Degradation | Xenobiotic degradation | Benzene | K16249+K16243+K16244+K16242+K16245+K16246 | M00548 |
| D080501 | D0805 | D08 | Degradation | Xenobiotic degradation | Benzoate | (K05549+K05550+K05784) K05783 | M00551 |
| D080502 | D0805 | D08 | Degradation | Xenobiotic degradation | Benzoate | K04116 K04117 K07534 K07535 K07536 | M00540 |
| D080601 | D0806 | D08 | Degradation | Xenobiotic degradation | Anthranilate | (K05599+K05600+K11311),(K16319+K16320+K18248+K18249) | M00637 |
| D080701 | D0807 | D08 | Degradation | Xenobiotic degradation | Catechol | K03381 K01856 K03464 (K01055,K14727) | M00568 |
| D080702 | D0807 | D08 | Degradation | Xenobiotic degradation | Catechol | (K00446,K07104) ((K10217 K01821 K01617),K10216) (K18364,K02554) (K18365,K01666) (K18366,K04073) | M00569 |
| D080801 | D0808 | D08 | Degradation | Xenobiotic degradation | Cumate | (K10619+K16303+K16304+K18227) K10620 K10621 K10622 K10623 | M00539 |
| D080901 | D0809 | D08 | Degradation | Xenobiotic degradation | Biphenyl | (K08689+K15750+K18087+K18088) K08690 K00462 K10222 | M00543 |
| D081001 | D0810 | D08 | Degradation | Xenobiotic degradation | Carbazole | K15751 (K15754+K15755) K15756 | M00544 |
| D081101 | D0811 | D08 | Degradation | Xenobiotic degradation | Benzoyl-CoA | ((K04112+K04113+K04114+K04115),(K19515+K19516)) K07537 K07538 K07539 | M00541 |
| D081201 | D0812 | D08 | Degradation | Xenobiotic degradation | Naphthalene | (K14579+K14580+K14578+K14581) K14582 K14583 K14584 K14585 K00152 | M00534 |
| D081301 | D0813 | D08 | Degradation | Xenobiotic degradation | Salicylate | K18242+K18243+K14578+K14581 | M00638 |
| D081401 | D0814 | D08 | Degradation | Xenobiotic degradation | Terephthalate | (K18074+K18075+K18077) K18076 | M00624 |
| D081501 | D0815 | D08 | Degradation | Xenobiotic degradation | Phthalate | (K18068+K18069) K18067 K04102 | M00623 |
| D081601 | D0816 | D08 | Degradation | Xenobiotic degradation | Phenylacetate | K01912 (K02609+K02610+K02611+K02612+K02613) K15866 K02618 K02615 K01692 K00074 | M00878 |
| D081701 | D0817 | D08 | Degradation | Xenobiotic degradation | Trans-cinnamate | (((K05708+K05709+K05710+K00529) K05711),K05712) K05713 K05714 K02554 K01666 K04073 | M00545 |
| D081801 | D0818 | D08 | Degradation | Xenobiotic degradation | Caffeine | K21722 K21723 K21724 | M00915 |
| D081901 | D0819 | D08 | Degradation | Xenobiotic degradation | Mercury | 4.99.1.2 1.16.1.1 | P641-PWY |
| D090101 | D0901 | D09 | Degradation | Antibiotic degradation | Penicillin | K18698,K18699,K18796,K18767,K18797,K19097,K19317,K18768,K18970,K19316,K22346,K18795,K19218,K19217,K17836,K18766 | NA |
| D090201 | D0902 | D09 | Degradation | Antibiotic degradation | Carbapenem | K17837,K18782,K18781,K18780,K19099,K19216 | NA |
| D090301 | D0903 | D09 | Degradation | Antibiotic degradation | Cephalosporin | K19095,K19096,K19100,K19101,K19214,K19215,K20319,K20320,K01467 | NA |
| D090401 | D0904 | D09 | Degradation | Antibiotic degradation | Oxacillin | K17838,K18790,K18791,K19098,K18792,K19213,K21276,K18793,K18971,K22352,K19209,K18976,K18973,K18794,K18972,K21277,K19210,K19211,K19212,K22335,K19319,K22331,K22351,K19320,K19318,K19321,K19322,K21266,K22334,K22333,K22332 | NA |
| D090501 | D0905 | D09 | Degradation | Antibiotic degradation | Streptogramin | K19349,K19350 | NA |
| D090601 | D0906 | D09 | Degradation | Antibiotic degradation | Fosfomycin | K21252 | NA |
| D090701 | D0907 | D09 | Degradation | Antibiotic degradation | Tetracycline | K08151,K08168,K18214,K18218,K18220,K18221 | NA |
| D090801 | D0908 | D09 | Degradation | Antibiotic degradation | Macrolide | K06979,K08217,K18230,K18231,K21251 | NA |
| D091001 | D0910 | D09 | Degradation | Antibiotic degradation | Chloramphenicol | K00638,K08160,K18552,K18553,K18554,K19271 | NA |
| D091101 | D0911 | D09 | Degradation | Antibiotic degradation | Lincosamide | K18236,K19349,K19350,K19545 | NA |
| D091201 | D0912 | D09 | Degradation | Antibiotic degradation | Streptothricin | K19273,K20816 | NA |
| S010101 | S0101 | S01 | Structure | Cellular structure | Peptidoglycan | 2.5.1.7 1.3.1.98 6.3.2.8 6.3.2.9 6.3.2.13 6.3.2.10 2.7.8.13 2.4.1.227 2.4.1.129 | PEPTIDOGLYCANSYN-PWY |
| S010102 | S0101 | S01 | Structure | Cellular structure | Peptidoglycan | 2.5.1.7 1.3.1.98 6.3.2.8 6.3.2.9 6.3.2.7 6.3.2.10 2.7.8.13 2.4.1.227 6.3.5.13 6.3.1.12 2.4.1.129 3.4.16.4 | PWY-6471 |
| S010103 | S0101 | S01 | Structure | Cellular structure | Peptidoglycan | 2.5.1.7 1.3.1.98 6.3.2.8 ((6.3.2.53 5.1.1.23),6.3.2.9) 6.3.2.13 6.3.2.4 6.3.2.10 2.7.1.66 2.7.8.13 2.4.1.227 | map00550 |
| S010301 | S0103 | S01 | Structure | Cellular structure | Teichoic acid | 3.6.1.27 2.7.8.33 2.4.1.187 2.7.8.44 ((2.7.8.12 2.4.1.52),(2.7.8.46 2.7.8.47 2.4.1.53),(2.7.8.45 2.7.8.14 (2.4.1.70,2.4.1.355))) 6.1.1.13 | map00552 |
| S010401 | S0104 | S01 | Structure | Cellular structure | Lipoteichoic acid | ((2.4.1.315 2.7.8.20),2.4.1.337) 6.1.1.13 | PWY-7817 |
| S010501 | S0105 | S01 | Structure | Cellular structure | Lipopolysaccharide | 2.3.1.129 3.5.1.108 2.3.1.191 2.4.1.182 2.7.1.130 2.4.99.12 2.4.99.13 2.3.1.241 2.3.1.243 | M00060 |
| S010502 | S0105 | S01 | Structure | Cellular structure | Lipopolysaccharide | K00677 K02535 K02536 (K03269,K09949) K00748 K00912 K02527 K02517 K02560 | M00060 |
| S010503 | S0105 | S01 | Structure | Cellular structure | Lipopolysaccharide | 5.3.1.28 2.7.1.167 3.1.3.82 2.7.7.70 5.1.3.20 | M00064 |
| S010504 | S0105 | S01 | Structure | Cellular structure | Lipopolysaccharide | K03271 (K03272,K21344) K03273 (K03272,K21345) K03274 | M00064 |
| S020101 | S0201 | S02 | Structure | Appendages | Flagellum | (K02411,K02412,K02413,K02418,K02419,K02420,K02421,K13820,K02400,K02401,K03516) (K02410,K02416,K02417) (K02409,K02394,K02386,K02393,K02415) (K02408,K02414,K02387,K02388,K02389,K02391,K02392,K02395,K02390,K02396,K02397,K02399) (K24343,K24344,K24344,K09860,K24346,K21217,K21218) (K18475,K02406,K02407,K02422,K02423) (K02403,K02402,K02405,K02556,K02557,K10564,K10565) | map02040 |
| S020102 | S0201 | S02 | Structure | Appendages | Flagellum | (K02411,K02412,K02413,K02418,K02419,K02420,K02421,K13820,K02400,K02401,K03516) (K02410,K02416,K02417) (K02409,K02394,K02386,K02393,K02415) (K02408,K02414,K02387,K02388,K02389,K02391,K02392,K02395,K02390,K02396,K02397,K02399) (K18475,K02406,K02407,K02422,K02423) (K02403,K02402,K02405,K02556,K02557,K10564,K10565) | map02040 |
| S020201 | S0202 | S02 | Structure | Appendages | Pilus | (K02666 K02656) (K02653 K02652 K02669) (K02662 K02663 K02664 K02665) K02650 (K02671 K02672 K02673 K02655) K02674 | NA |
| S020202 | S0202 | S02 | Structure | Appendages | Pilus | K07350 (K07349 K07348) K07345 | NA |
| S020203 | S0202 | S02 | Structure | Appendages | Pilus | K12522 (K12521 K12520 K12523) K12517 | NA |
| S030101 | S0301 | S03 | Structure | Spore | Spore | (K07699 K03091) (K06378 K06379 K06382 K06383 K06387) (K06374 K06390 K06392 K06393 K06394 K06396 K06397 K06398 K06399 K06405 K06407 K06409 K04769 K06438) | NA |
